# Harnessing homeostatically active RhoC at cell junctions preserves human endothelial barrier function during inflammation

**DOI:** 10.1101/2024.05.17.594667

**Authors:** Natalia Colás-Algora, Pablo Muñoz-Pinillos, Susana Barroso, Cristina Cacho-Navas, Álvaro Caballero, Gema Cerro-Tello, Gema de Rivas, Martín González-Fernández, Ignacio Jiménez-Alfaro, Manuel Fresno, Catalina Ribas, Alberto Paradela, Eduardo López-Collazo, José Jesús Fernández, Jaime Millán

## Abstract

Rho GTPases are molecular targets of bacterial toxins that modulate their enzymatic activity. RhoA, RhoB and RhoC are almost identical and play critical roles in generating actomyosin-mediated contractile forces that cause endothelial hyperpermeability during inflammation. Searching for new treatments to modulate endothelial integrity, we demonstrate that the specific and simultaneous activation of these three Rho GTPases with a chimeric recombinant toxin does not induce cell contraction but enhances homeostatic endothelial barrier function, increases reticular adherens junctions and preserves the microvascular endothelium in response to pathological inflammatory challenges *in vitro* and *in vivo*. This pro-barrier effect is specifically mediated by RhoC, whose activity is increased by cell confluence. The uniqueness of RhoC relies on an arginine 188 within its hypervariable region that determines its junctional localization, high homeostatic activity, and barrier-protective function. Quantitative proteomics revealed that RhoC regulates the expression of myosin light chain proteins and junction-stabilizing actomyosin. Thus, harnessing the activity of RhoC represents a potential therapy for strengthening endothelial barriers during pathological inflammation.

## INTRODUCTION

Systemic and exacerbated inflammatory responses, such as those caused by infection or trauma in sepsis, leads to life-threatening organ edema and dysfunction. The uncontrolled secretion of inflammatory mediators in these pathologies generates a cytokine storm that enters the circulation and alters endothelial permeability, increasing vascular leakage even if the causes giving rise to the damaging inflammatory response are successfully identified and treated ^1,2^. Current strategies for preventing fatal tissue edema during sepsis are focused on inhibiting this inflammatory signaling and stabilizing the vascular tone ^2–4^. However, given the importance of the vasculature in this pathology ^1,5^, a complementary strategy, which has yet to be thoroughly explored, may consist of transiently reinforcing the endothelium during the critical illness period, regardless of the specific mediators responsible for vascular leakage.

Endothelial integrity is regulated by the family of Rho GTPases, molecular switches that control multiple functions, including the actin cytoskeleton, a major endothelial scaffold. In the current paradigm of molecular pathways governing endothelial barrier function, RhoA, the founder member of this family, is the central regulator of stress fiber formation and F-actin-mediated contraction, which compromise the endothelial integrity in response to acute inflammatory signals such as thrombin or histamine both *in vivo* and *in vitro* ^6,7^. This GTPase acts in opposition to the GTPase Rac1, which mediate endothelial barrier enhancement ^8–10^. RhoA belongs to a subfamily that comprises another two GTPases, RhoB and RhoC, which share 88% amino acid identity ^11^. They regulate common effectors, such as Rho kinases (ROCK) and play redundant, complementary and specific roles depending on the cellular context ^10–12^. The main sequence divergence between RhoA, RhoB, and RhoC is localized in the C-terminal hypervariable region, which is essential for their prenylation and membrane localization ^13^.

Vascular endothelial (VE)-cadherin-dependent adherens junctions (AJs) are the main regulators of endothelial barrier integrity ^14^. VE-cadherin partners with cytoplasmic catenins, which connect AJs to the underlying actin cytoskeleton that stabilizes this junctional protein complex. VE-cadherin cross talks with other junctional protein complexes, such as those formed by PECAM-1, into distinct junctional structures with reticular morphology that stabilize contacts between cells ^15,16^. PECAM-1 and VE-cadherin also form a mechanosignaling module ^17^ that signals to actin stress fibers through Rho GTPases. Surface VE-cadherin also participates in cell survival, collective endothelial cell migration and gene transcription ^14,18–20^ so its expression is closely regulated even when endothelial barrier integrity is transiently disrupted by inflammatory mediators ^21^.

We had previously reported that RhoB has a specific function as a negative regulator of barrier function in an inflammatory context in endothelial cells ^10^ and, as had other laboratories, that inhibiting the RhoA subfamily effector ROCK prevents acute contraction induced by inflammatory mediators junctions ^15,22,23^. Here, unexpectedly, we found that activating endogenous RhoA, RhoB and RhoC with a chimeric protein containing the catalytic domain of the cytotoxic necrotizing factor y (CNFy) toxin from *Yersinia pseudotuberculosis*, far from inducing contraction, increases reticular AJs and perijunctional F-actin, maintains endothelial barrier function *in vitro*, prevents local microvascular disruption in response to inflammatory mediators *in vivo*, and reduces organ edema upon systemic LPS stimulation. Hence, this RhoA activator emerges as a potential therapeutic approach for pathologies associated with endothelial barrier dysfunction. More importantly, these findings have led us to demonstrate that, despite its similarity to RhoA and RhoB, RhoC plays an opposite, barrier-protective role that mediates the effect of this chimeric protein. One single, positively charged, C-terminal R188 residue in RhoC, which is not present in the equivalent position of RhoA or RhoB, determines the localization, high activity and pro-barrier function of this GTPase. Hence, the spatiotemporal control of the RhoA GTPase subfamily activity and the relative expression of each of its three members determine its pro-barrier or anti-barrier roles, which could be therapeutically exploited.

## RESULTS

### Activation of endogenous RhoA, RhoB and RhoC by CN03 increases reticular AJs and strengthens endothelial barrier function

Cytotoxic necrotizing factors (CNFs) are bacterial toxins that induce deamidation of a glutamine residue in the Switch II region of Rho GTPases, blocking their GTPase activity and thus constitutively activating these enzymes. Some CNFs, such as CNF1, induce deamidation of various Rho GTPases, such as those from the Rac, RhoA and Cdc42 subfamilies. Some others, such as CNFy, are specific to the RhoA subfamily ^24–26^. We had previously reported an essential role of RhoA and RhoB decreasing human endothelial barrier function in an inflammatory context ^10,15^. To investigate further the role of the RhoA subfamily of GTPases in human endothelial cells, we used CN03, a commercially available recombinant protein that contains the catalytic domain of a CNF toxin and that is conjugated to a cell-penetrating peptide that translocates the toxin into the cell after 2-3 hours of exposure ^27^. CN03 activated RhoA, RhoB and RhoC but not Cdc42 or Rac1 in primary human vascular endothelial cells (Fig. 1a and Extended Data Fig. 1a) confirming our previous analyses in other cell types ^27^. Interestingly, CN03 promoted the formation of robust VE-cadherin-mediated AJs with a reticular morphology in various endothelial beds such as human umbilical vein endothelial cells (HUVECs) (Fig. 1b, boxed areas) and human dermal microvascular endothelial cells (HDMVECs) (Fig. 1c, boxed areas). Reticular AJs are cell-cell contact domains where VE-cadherin crosstalk with the junctional adhesion receptor PECAM-1 ^15^. The reticular architecture of VE-cadherin and PECAM-1 reduces endothelial permeability, has been found *in vivo* and *in vitro* ^4,15,28,29^ and is disrupted by inflammatory cytokines, anti-PECAM-1 blocking antibodies and autophagy ^15,28^. Indeed, VE-cadherin and PECAM-1 staining intensities at cell–cell contacts, as well as areas covered by reticular AJs, were notably higher in cells exposed to CN03 (Fig. 1b, c), which was linked to an increase of total and surface protein expression of VE-cadherin (Fig. 1d, e). In contrast, ZO-1, which forms endothelial tight junctions, was not found in reticular junctions and did not significantly change expression in response to CN03 exposure (Extended Data Fig. 1b, c). To investigate further the effect of CN03 on VE-cadherin-mediated AJs, we generated a construct to express a photoconvertible VE-cadherin-Dendra2 chimera in endothelial cells. Dendra2 is a fluorescent protein that undergoes an emission shift from 488-nm to 543-nm in response to irradiation with UV light, which allows to analyze protein diffusion in the photoconverted regions by time-lapse confocal microscopy. Confluent cells were exposed or not to CN03 for 3 h before photoconversion (Fig. 1f). Photoconverted VE-cadherin-Dendra2 at reticular AJs exhibited lower diffusion rate than when this junctional protein was distributed with linear morphology, indicating that the reticular disposition favors the stability of AJs. In fact, CN03 not only increased reticular AJs but also reduced the dispersion of VE-cadherin, especially when this junctional protein was distributed in a reticular pattern (Fig.1f, bottom graphs).

**Figure 1.**
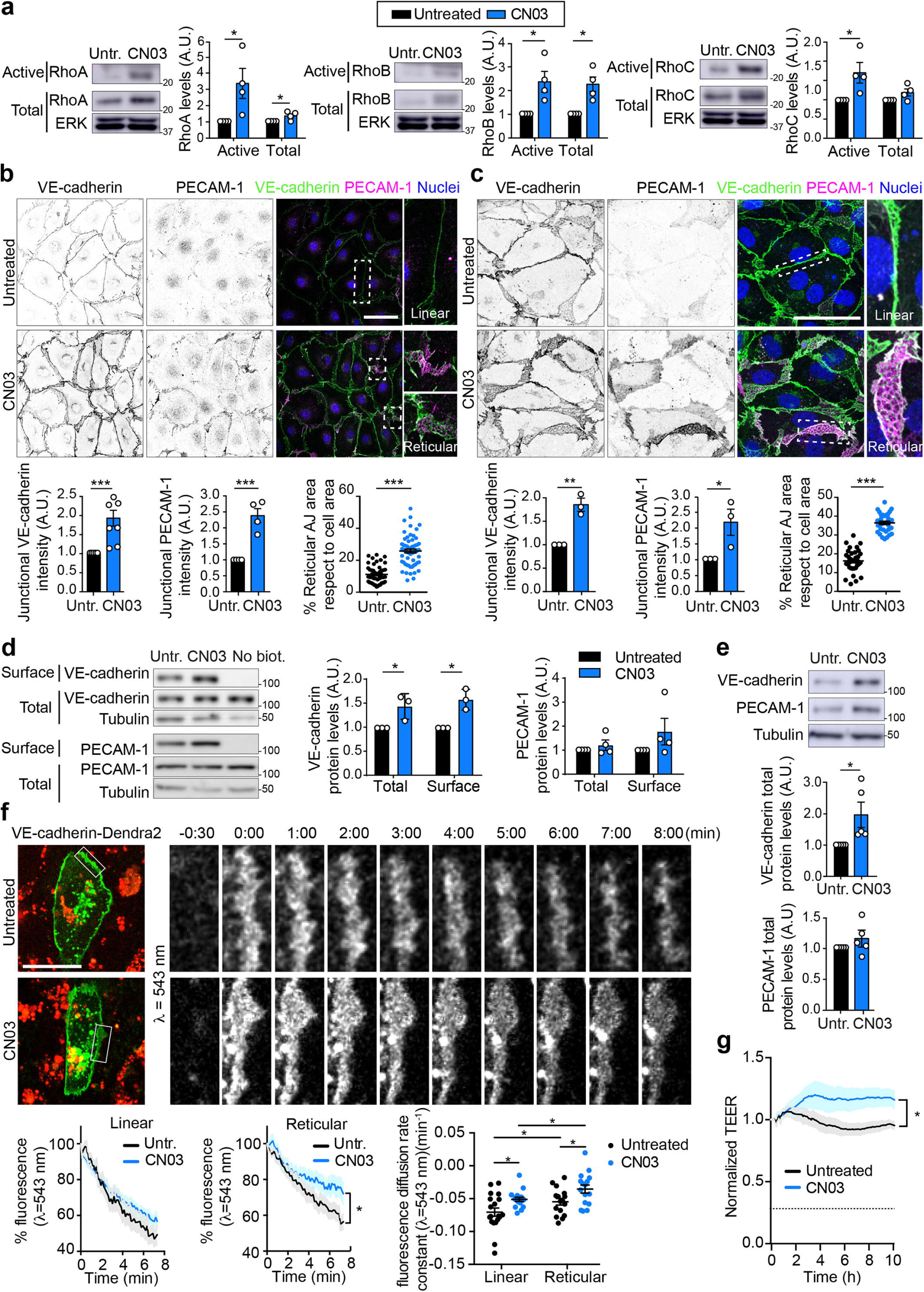
CN03 activates RhoA, RhoB and RhoC, increases reticular AJs and reinforces endothelial barrier function. **a,** Confluent HUVECs were exposed or not to 1.5 μg/ml CN03 for 3 h, lysed and the indicated active GTPases were detected by pull-down assays with a recombinant protein containing the Rho binding domain (RBD) of Rhotekin conjugated to GST. **b,c,** Confluent HUVECs **(b)** or HDMVECs **(c)** were exposed to 1.5 μg/ml or 5 μg/ml CN03, respectively, for 3 h, fixed and stained for VE-cadherin and PECAM-1. Right images show two-fold enlargements of the squared areas highlighting the morphology of linear (top) and reticular (bottom) AJs. Bottom plots show quantification of junctional VE-cadherin and PECAM-1 levels and the percentage of reticular AJs area. Nuclei were stained with DAPI. Scale bars, 50 μm. **d,** CN03 increases total and surface protein expression of VE-cadherin in HUVECs. Surface-biotinylated proteins were isolated by pull-down assay with NeutrAvidin-agarose in HUVECs exposed or not to CN03 as in **(a)**. VE-cadherin and PECAM-1 protein levels from the surface pull-down fraction were compared with total protein levels from the lysates by western blot. Non-biotinylated cells (No biot.) were analyzed in parallel as a control of the pull-down assay. Blots of tubulin are shown as loading controls. Plots show the mean□±□SEM from at least three independent experiments. **e,** Protein expression changes of PECAM-1 and VE-cadherin in HDMVEC exposed to CN03 as in **(b)**. Plots show the mean□±□SEM from five independent experiments. **f,** CN03 stabilizes VE-cadherin preferentially at reticular AJs. HUVECs expressing photoconvertible VE-cadherin-Dendra2 were treated with 1.5 μg/ml CN03 for 3h, then photoconverted at cell-cell junctions and subjected to time-lapse confocal analysis for 8 min exciting at 543 nm of wawelength (λ). Scale bar, 30 μm. Bottom graphs. Fluorescence decrease in the photoconverted area was calculated as percentage of maximum fluorescence detected at time 0 (left and central graphs) and from these values the constant of fluorescence reduction was calculated (right graph). In accordance with the morphological distribution of VE-cadherin-Dendra2, regions were classified as linear or reticular. At least 12 cells per condition were photoconverted in 3 independent experiments. **g,** TEER analysis with an ECIS system of HDMVEC exposed to CN03 as in **(c)**. *, P< 0.05; **, P< 0.01; *** P<0.001.

To analyze in a quantitative manner whether the increase of reticular AJs affects endothelial barrier integrity, we measured multifrequency transendothelial resistance (TEER) of primary human endothelial cells exposed or not to CN03, using an ECIS system. CN03 increased TEER, which confirmed that this RhoA subfamily activator reinforces endothelial barrier integrity (Fig. 1g). Moreover, ECIS can mathematically model multifrequency TEER measurements and separate it into three parameters of different biological meanings: Rb, which provides the contribution of paracellular permeability and cell-cell junctions to barrier integrity; α, which measures cell-matrix interactions; and Cm, which is relative to the membrane capacitance ^4,30,31^. The significant increase of TEER observed in cells treated with CN03 was mostly caused by changes in Rb values, but not in α or Cm, indicating that this recombinant protein exerts a protective effect related with junction-related paracellular permeability (Extended Data Fig. 1d).

To elucidate why this reticular disposition strengthens endothelial barrier integrity, we explored the structure of reticular AJs in greater depth. Reticular areas were first localized for analysis by transmission electron microscopy (EM) by expressing PECAM-1-GFP (not shown). We then acquired EM micrographs of serial sections (Fig. 2a) and performed a 3D reconstruction (Fig. 2b). The 3D analyses revealed superimposed plasma membranes between adjacent cells in which their interface was irregular and featured invaginations that suggested active processes of internalization and intercellular communication (Fig. 2b, arrowheads). These results indicate that reticular AJs provide an extra source of plasma membrane that increases the surface area in which to form intercellular protein complexes. The extension of these cellular interfaces potentially helps to maintain the integrity of the endothelial monolayer by coping with changes in vessel diameter and cellular shape caused by the mechanical stress induced by inflammation, blood pressure, trauma, etc. Thus, collectively, our findings indicate that CN03 strengthens endothelial cell-cell contacts by increasing PECAM-1 and VE-cadherin-rich reticular AJs, which extend these complexes into junctional overlapping areas, thereby reducing homeostatic paracellular permeability in the human endothelium.

**Figure 2.**
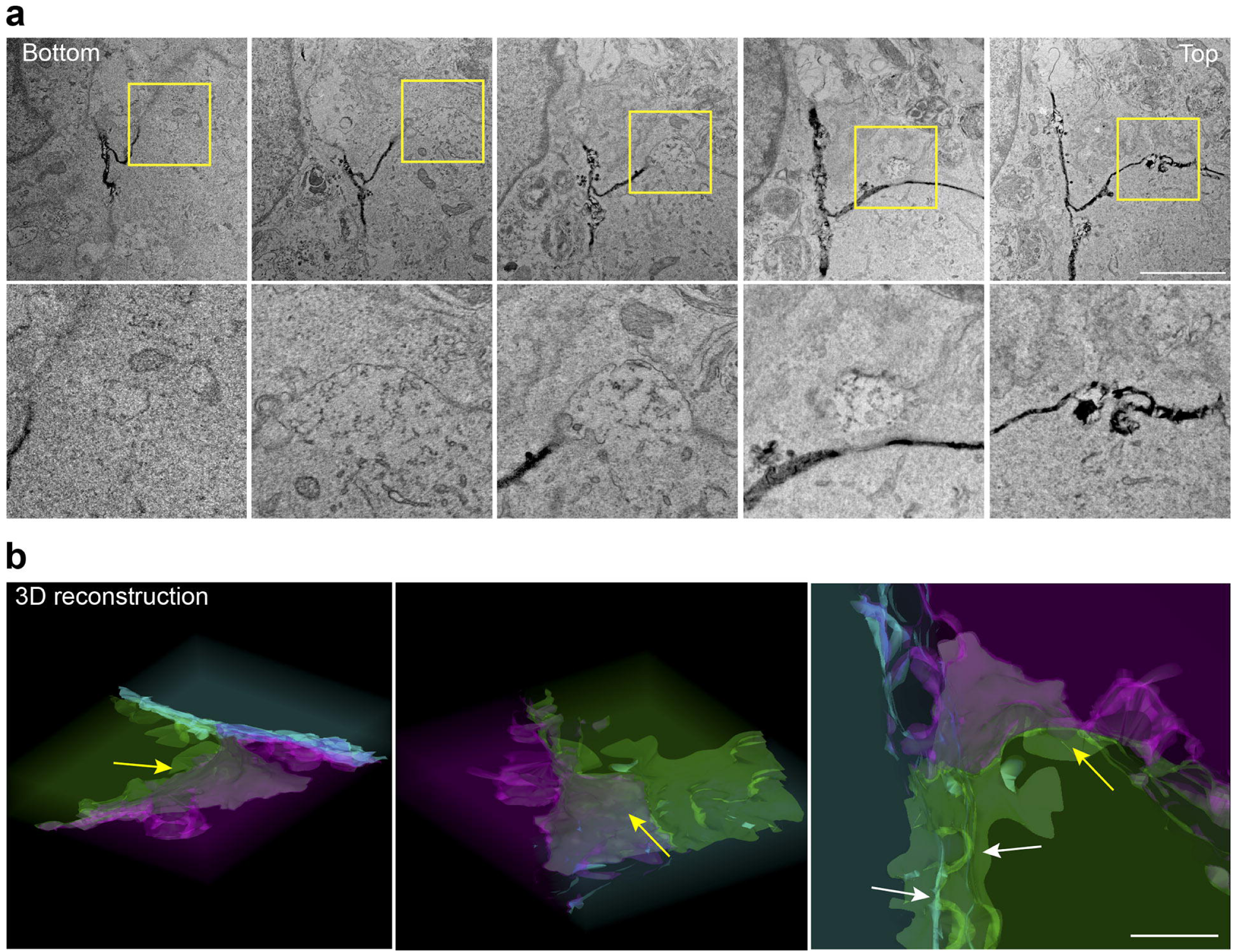
Reticular AJs are formed at overlapping junctional areas. **a,** Top images show representative micrographs of 27 consecutive transmission electron microscopy sections of microvascular endothelial cells. Bottom images are three-fold enlargements of the boxed areas showing membrane invaginations. **b,** 3D reconstruction of these sections reveals that reticular junctions are formed by overlapping junctional plasma membranes enriched in intertwined invaginations. In the model, plasma membranes of different cells are delineated in different colors and their cytoplasms in semi-transparent textures. White and yellow arrows point at invaginations detected between two different overlapping cells. Scale bar in electron microscopy micrographs, 5 μm. Scale bar in the 3D reconstruction, 2 μm.

Given the unexpected reinforcement of human endothelial barrier by CN03, as a final control, we identified the exact nature of CN03 by mass spectrometry analysis. This revealed that it is a chimeric toxin containing the N-terminal domain of CNF1 from *Escherichia coli* and the C-term catalytic domain of CNFy from *Yersinia pseudotuberculosis,* which specifically activates RhoA subfamily proteins (Extended Data Fig. 1e). This fusion protein is very similar to that previously reported by Hoffman et al. ^32^. In addition, to address the role of the RhoA subfamily activity using an alternative strategy, we analyzed the effect of inhibiting these three GTPases with C3 transferase, an exoenzyme from *Clostridium botulinum* ^10,33^. C3 transferase did not significantly change the TEER values. However, multifrequency TEER modeling revealed a massive decrease in Rb, concomitant with an increase in α and Cm, indicating that, despite not altering the TEER values, C3 induces a substantial increase in paracellular permeability that is compensated by an increase in adhesion to the substratum and changes in the membrane capacitance (Extended Data Fig. 2a). Consistent with these results, confocal microscopy analyses revealed that C3, which massively reduced F-actin staining, induced many intercellular gaps in confluent cells (Extended Data Fig. 2b, arrowheads). Overall, the use of different recombinant proteins with capacity to modulate the activity of endogenous RhoA, RhoB and RhoC, indicate that the activity of the RhoA subfamily is necessary for maintaining the barrier properties of primary human endothelial cells.

### CN03 ameliorates endothelial barrier disruption *in vitro,* and local and systemic vascular leakage *in vivo,* in response to inflammatory challenges

We then analyzed the effect of CN03 on endothelial cells exposed to inflammatory challenges that induce pathological hyperpermeability. Bacterial lipopolysaccharide (LPS) is a major activator of the innate immune system during infection and is used as a model to induce endothelial barrier disruption *in vitro* and to recapitulate acute sepsis *in vivo* ^34^. LPS caused a significant reduction of TEER and Rb values in microvascular endothelial cells (Fig. 3a). CN03 significantly attenuated such decrease, recovering these values to levels close to those before to the exposure to the inflammatory challenge (Fig. 3a). CN03 increased VE-cadherin and PECAM-1 at junctions in the presence of LPS, reducing their dispersion, and decreased the formation of intercellular gaps (Fig. 3b). During sepsis, circulating blood with high levels of inflammatory mediators induces microvascular leakage and edema. The lung is one of the main organs affected by microvascular hyperpermeability. To mimic this systemic pathology in human cells, we analyzed the effect of two sets of sera, collected from healthy donors and severely ill patients with sepsis, on lung microvascular endothelial cell barriers (HLMVECs), as previously described ^4^. Septic sera induced a stronger disruption of the HLMVEC barrier than sera from healthy donors (Fig. 3c). Interestingly, septic sera also induced cell contraction and the appearance of massive intercellular gaps that were significantly reduced by CN03 (Fig. 3d,e). Hence, overall, reinforcing the human endothelium with CN03, strongly protects these cellular barriers from pathological proinflammatory challenges.

**Figure 3.**
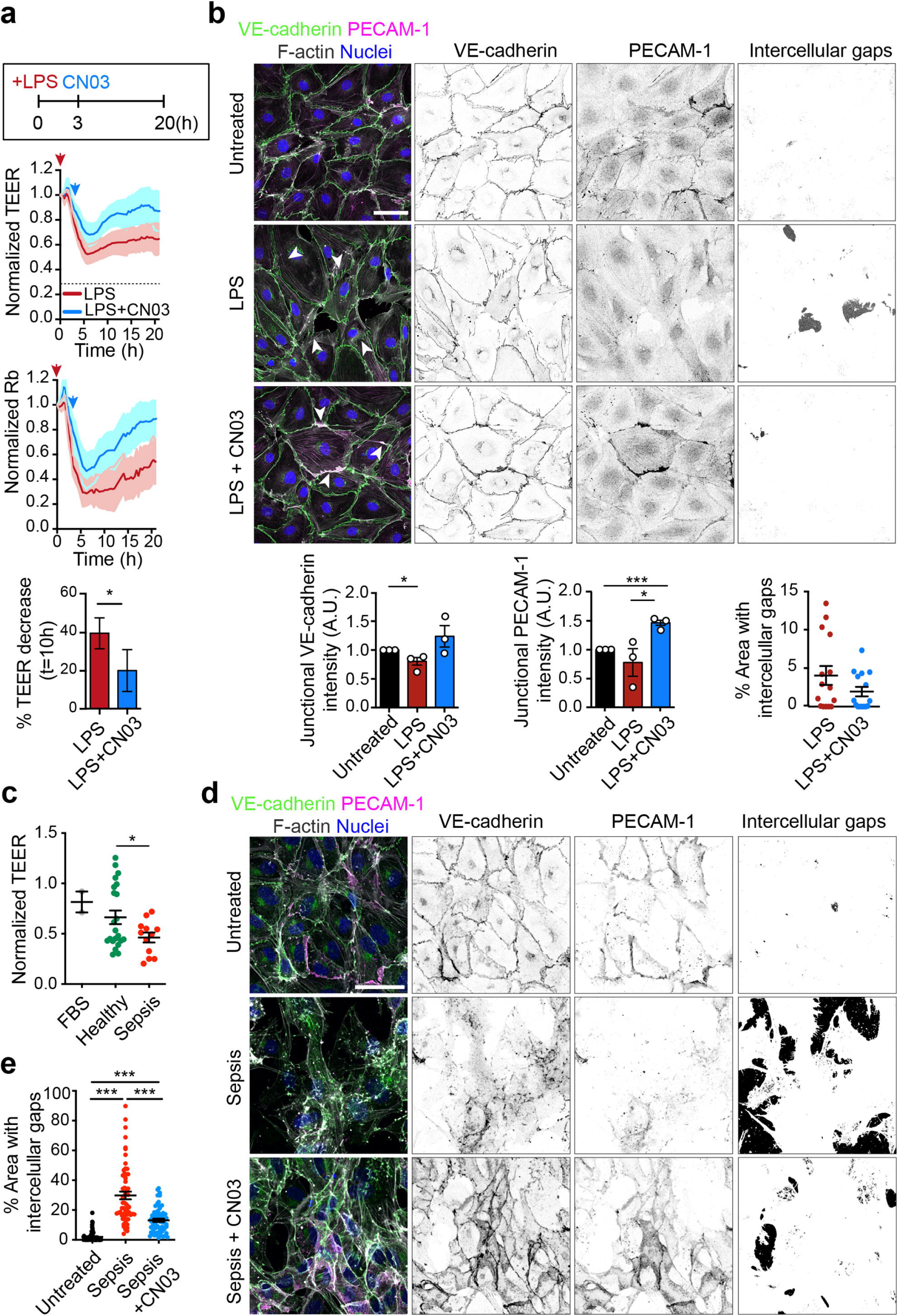
CN03 protects human endothelial barrier from pathological inflammatory stimuli. **a,** TEER and Rb measurements of confluent HDMVECs exposed to bacterial LPS and 3 h later treated or not with 5 μg/ml CN03. Bottom plots show the quantification of TEER decrease after 10 h of LPS stimulation. **b,** Confluent HDMVEC were incubated with LPS and CN03 as in **(a)**, fixed and stained for VE-cadherin, PECAM-1 and F-actin. Bottom left and central plots show the quantification of junctional VE-cadherin and PECAM-1 levels. Bottom right plot shows the quantification of intercellular gaps identified by automatic image processing with Fiji. **c,** Confluent HLMVEC were exposed to 30% sera from healthy donors or sera from patients suffering severe sepsis and TEER was measured with an ECIS system for 48 h. 30% FBS was used as control. **d,e**, Endothelial cells were exposed or not to septic sera with or without CN03 for 24 h, fixed and stained for VE-cadherin, PECAM-1 and F-actin. Intercellular gaps were identified by automatic image processing with Fiji and quantified in **(e)**. Nuclei were stained with DAPI. Plots show the mean□±□SEM of cell quantifications from cells from three independent experiments. Scale bars, 50 μm. *, P< 0.05; *** P<0.001.

We then tested the effect of intravenous injection of CN03 in a mouse model of local induction of vascular leakage detected by Miles assay ^35^. Intravenous injection of Evans blue was followed by injection of LPS, VEGF and control saline buffer into the skin (Fig. 4a). CN03 pretreatment significantly reduced vascular leakage by almost 50% in LPS-injected skin patches, and had also a milder effect on VEGF-induced vascular leakage. In contrast, CN03 did not affect homeostatic vascular leakage in liver, lung or kidney (Extended Data Fig. 3a). Systemic bacterial, viral or fungal infection causes a cytokine storm that triggers endothelial barrier disruption, organ edema and failure during sepsis ^1^. To evaluate the effect of CN03 systemically *in vivo*, the acute phase of bacterial sepsis was mimicked by intraperitoneal injection of LPS in the presence of saline or CN03. CN03 significantly reduced the increase of TNF and IL-6 concentration in blood (Fig. 4b) as well as systemic vascular leakiness in livers, lungs and kidneys (Fig. 4c). In contrast, CN03 had no significant effect on LPS-induced hypoglycemia (Extended Data Fig. 3b). Therefore, our results clearly show that the activation of endogenous RhoA, RhoB and RhoC GTPases by CN03 reduces blood inflammatory cytokines, endothelial barrier disruption and vascular leakage when systemic inflammation is induced *in vivo*.

**Figure 4.**
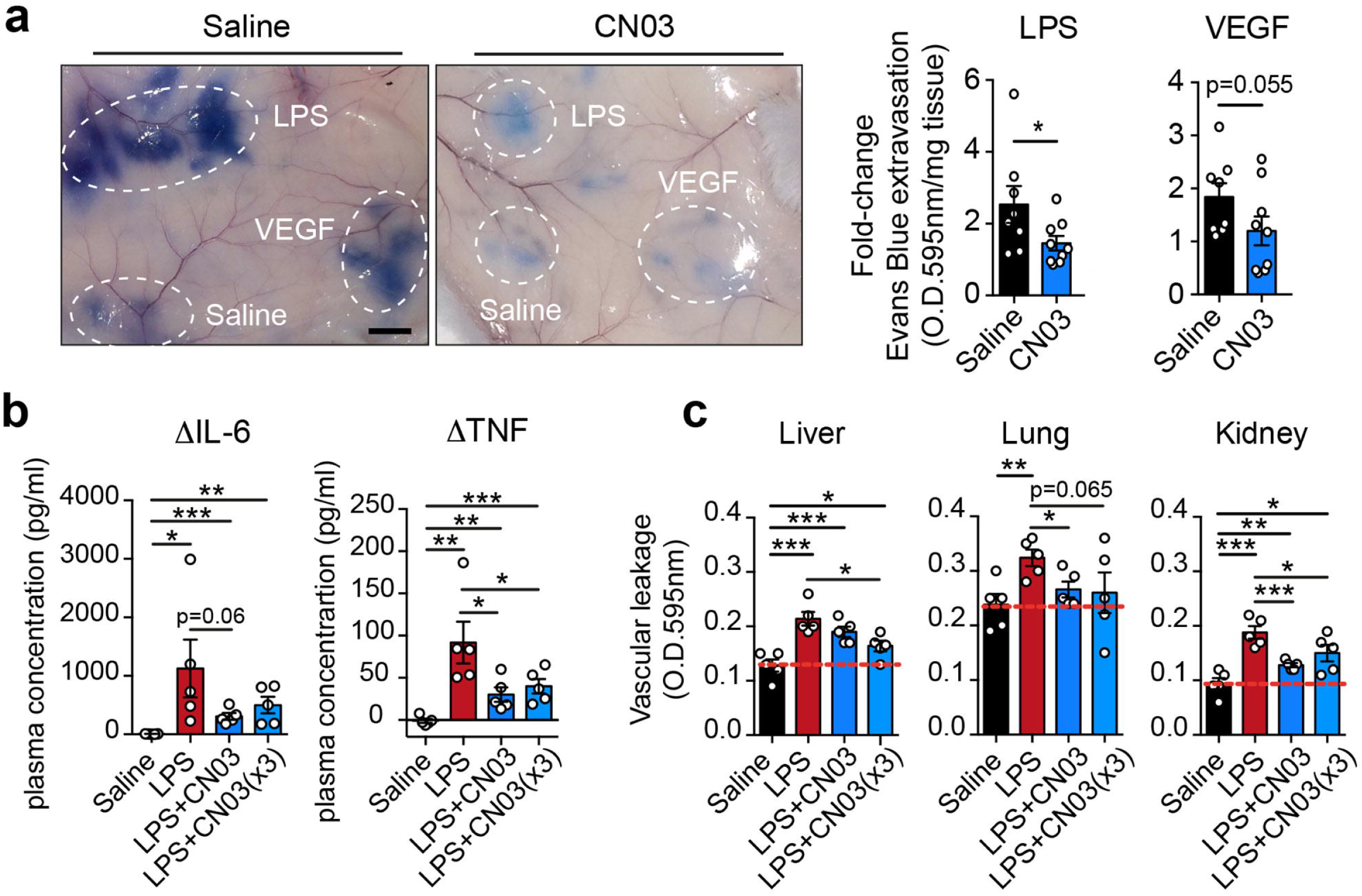
CN03 reduces local and systemic edema and circulating cytokines in response to LPS injection. **a,** Representative images of mouse skin locally injected with 20 μg LPS, 50 ng VEGF, or saline serum, 4 h after intra-peritoneal injection of saline serum or 0.175 mg/kg CN03, and 15 min after retro-orbital injection of Evans Blue (EB). Right plots. EB was extracted from skin patches and quantified relative to values of saline-injected patches (right plots). Nine animals per condition were used. **b,c,** Blood was initially collected for cytokine and glucose analyses. Mice were then intra-peritoneally injected with saline, LPS+saline, LPS + 0.175 mg/kg CN03 and LPS + 0.525 mg/kg (CN03×3) for 3.5 h, EB was retro-orbitally injected for 15 min and blood was collected again to the measured IL-6 and TNF-α in plasma **(b)**. The graphs show the differences (Δ) in cytokine concentration in blood measured after the injection of LPS, compared to the values obtained before the injection, for each of the mice. Animals were then sacrificed and vascular leakage was measured by quantifying optical density (O.D.) of EB at 595 nm in 100 mg of the indicated tissues **(c)**. Dotted line shows mean values of mice intra-peritoneally injected with saline serum. Plots show the mean□±□SEM. Five animals per condition were used. *, P< 0.05; **, P< 0.01; *** P<0.001.

### The effect of CN03 on endothelial barrier function depends on RhoC expression

To gain insight into the contribution of each RhoA subfamily member to the effect of CN03 on endothelial barrier function, we compared the morphological changes induced by this chimeric protein on cell-cell junctions of endothelial cells silenced, or not, for each of the three *RHOA* subfamily genes. First, single *RHO* gene knockdown had a significant effect on the expression of the other family members in HUVECs (Fig. 5a), as previously reported ^10^, and in HDMVECs (Extended Data Fig. 4a). Such upregulation also involved an increase in the activity of the corresponding Rho protein, except for RhoB, which remained mostly inactive (Extended Data Fig. 4b). These compensations suggest that the individual members of the RhoA subfamily of GTPases play additive and redundant roles in human endothelial cells. In contrast, single or double knockdown of these three GTPases did not significantly affect the expression or activity of Rac1 and had a moderate effect on the activity of Cdc42 (Extended Data Fig. 4c). ECIS analyses of single RHO gene silencing showed that RhoA depletion increased endothelial TEER and Rb, indicating that RhoA expression reduces barrier integrity and increases paracellular permeability. In contrast, RhoC depletion reduced TEER and Rb, suggesting that RhoC is necessary for maintaining barrier function and low paracellular permeability (Fig. 5b). No major changes were detected in other parameters related with endothelial barrier function (Extended Data Fig. 4d). In contrast with previous reports suggesting a significant role for RhoB in endothelial permeability ^36^, we observed no major effect on permeability when RhoB expression was reduced in HUVECs. This is consistent with previous data showing low levels of RhoB expression in HUVECs that had not been exposed to inflammatory mediators for several hours ^10^ and is also in accordance with the low levels of activation detected for this GTPase (Extended Data Fig. 4b). In addition, RhoC knockdown also resulted in a specific reduction of TEER and Rb in HDMVECs, indicating that proper RhoC expression is required for maintaining paracellular permeability in the microvascular endothelium as well (Extended Data Fig. 4e). Moreover, RhoA and RhoB knockdown did not affect the CN03-dependent increase of VE-cadherin and PECAM-1 at the cell periphery, whereas RhoC knockdown significantly disrupted the formation of cell-cell junctions (Fig. 5c, d). Cells lacking RhoC also failed to localize VE-cadherin and PECAM-1 to cell-cell contacts upon exposure to CN03, despite being initially seeded at high density to promote confluence (Fig. 5c, d and Extended Data Fig. 4f). Furthermore, *RHOC* gene silencing not only dispersed VE-cadherin and PECAM-1 but also markedly reduced the expression of the former, as revealed by western blot analyses of cells transfected for four different siRNA oligonucleotides (Extended Data Fig. 5a). qPCR assays indicated that this reduction was not transcriptional (Extended Data Fig. 5b), but was related to protein stability, as RhoC knockdown reduced VE-cadherin half-life (Extended Data Fig. 5c). Finally, the three RhoA subfamily members were individually or simultaneously silenced, and the expression analyses were extended to additional components of cell-cell junctions. Although RhoA and RhoB moderately regulated the expression of some junctional proteins, and triple Rho silencing reduced the expression of other endothelial cadherins such as N-cadherin (Extended Data Fig. 5d, e) ^37^, the presence of RhoC was mostly necessary to maintain the expression of VE-cadherin and α-catenin. Collectively, these findings demonstrated that RhoC is the sole member of the subfamily that strengthens endothelial barrier function. Such enhancement occurs specifically through the stabilization of VE-cadherin protein and VE-cadherin-mediated AJs.

**Figure 5.**
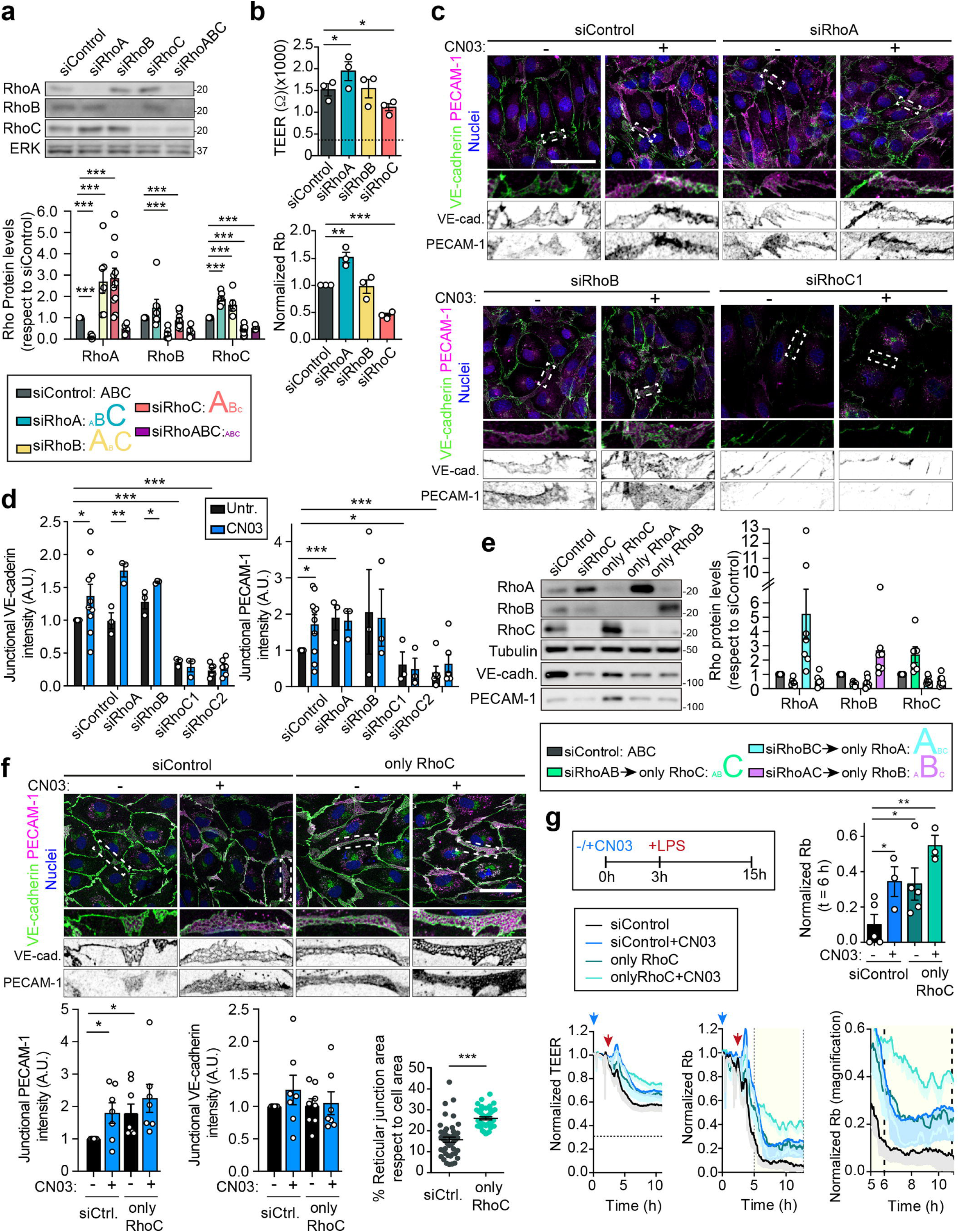
RhoC mediates the protective effect of CN03 on endothelial barrier function. **a,** Compensatory changes of expression of RhoA subfamily members upon single knockdown of RhoA, RhoB and RhoC. **b,** TEER and normalized Rb values of HUVECs transfected with the indicated siRNA for 72 h. **c,** Confocal analyses of HUVECs transfected with the indicated siRNA for 72 h, exposed to CN03 or not for 3 h, fixed and stained for VE-cadherin, PECAM-1 and F-actin. Bottom images are three-fold enlargements of the boxed areas. **d,** Relative quantification of VE-cadherin and PECAM-1 intensity at junctions. Includes the quantification of the effect of siRhoC2, shown in Extended Data Fig. 4f. **e,** Compensatory expression changes of the remaining RhoA subfamily member after double knockdown of the other two members. Double knockdowns generate cells expressing very high levels of only one RhoA subfamily GTPase (only RhoA, only RhoB, only RhoC). **f,** Increased junctional PECAM-1 and reticular AJs in only RhoC cells with respect to siControl cells. Cells were stained as in **(c)**. Bottom images are three-fold enlargements of the boxed areas. **g,** Reduced hyperpermeability to LPS, measured as the Rb value from TEER measurements, in only RhoC cells, which are more sensitive to the protective effect of CN03. Nuclei were stained with DAPI. Plots show the mean□±□SEM from at least three independent experiments. *, P< 0.05; **, P< 0.01; *** P<0.001.

To investigate the differences among the three members of the RhoA subfamily in greater detail, we analyzed the barrier properties of cells manipulated so that they predominantly expressed only one of these GTPase. Similar to the depletion of a single RhoA subfamily member, simultaneous depletion of two GTPases induced an upregulation of the remaining Rho by two-to five-fold, resulting in an overall enrichment of five-to ten-fold relative to the silenced GTPases in HUVECs (Fig. 5e). The double-depleted cells were thus termed “only RhoA”, “only RhoB” and “only RhoC”. Only RhoC cells did not alter the junctional recruitment of VE-cadherin (Fig. 5f) but did remarkably increased PECAM-1 expression (Fig. 5e) and junctional localization, and displayed abundant reticular, PECAM-1-enriched AJs (Fig. 5f), consistent with previous reports that attributed this particular junctional morphology to this adhesion receptor ^15^. Although only RhoA and only RhoC cells were able to maintain homeostatic barrier function, only RhoB cells showed reduced TEER and Rb values, indicating that the upregulation of this Rho GTPases cannot fully compensate the lack of the other two, probably due to its partial and specific endosomal localization (Extended Data Fig. 6a). Moreover, in contrast to only RhoC cells (Fig. 5f), only RhoA and only RhoB cells could failed to translocate both PECAM-1 and VE-cadherin to cell-cell junctions in response to the CN03 treatment (Extended Data Fig. 6b, c). Finally, only RhoC cells displayed reduced barrier hyperpermeability from 6 h onwards after LPS stimulation, which was comparable to the protective effect of CN03 (Fig. 5g and Extended Data Fig. 6d). Together, these experiments demonstrate that the RhoA subfamily is essential for regulating paracellular permeability, but that its pro- or anti-barrier function in the human endothelium depends on the relative expression of individual members of the family. Endothelial cell barriers that predominantly express RhoC are more resistant to inflammatory disruptors in a manner comparable to that observed upon CN03 exposure.

### RhoC regulates the expression of VE-cadherin by controlling perijunctional actomyosin through MLCK

To better understand the molecular mechanisms underlying the specific role of RhoC in maintaining cell-cell contacts and VE-cadherin protein stability, we performed label-free differential proteomics analyses between cells transfected with siRNA control and cells silenced for RhoC with two different siRNA oligonucleotides. In addition to detecting lower levels of some proteins forming AJs, already observed in the junctional protein screening (Extended Data Fig. 5e), such as α-catenin (Extended Data Table 1), quantitative proteomics revealed significant alterations in the expression of non-muscle myosin-II components (Fig. 6a-c). Non-muscle myosin II is an actin-based molecular motor forming a hetero-multimeric protein complex comprising homodimers of motor myosin heavy chains (MHCs) associated with two sets of pairs of myosin light chains (MLCs): regulatory and essential. The regulatory MLCs control ATP-dependent conformational changes in MHCs, which mediate contraction by sliding actomyosin filaments alongside each other in an antiparallel way ^38,39^. In addition, MLCK and Rho Kinase (ROCK) regulate myosin-II-mediated contraction by phosphorylating these regulatory MLCs. Quantitative proteomics revealed a consistent decrease of proteins related to myosin-II activity in RhoC depleted cells, such as the regulatory MLCs, MYL9 and MYL12B, and the essential MLCs, MYL6 and MYL6B (Fig. 6b) (Extended Data Table 1) ^40^. Western blot against MLC-2 (MYL9) and phospho-(S-19) MLC (MYL9 and MYL12B) ^41^ confirmed these changes and revealed that each of the three RhoA subfamily members contribute to the expression and phosphorylation of MLC, although RhoC silencing caused the strongest reduction (Fig. 6c). However, this additive effect on regulatory MLC did not explain de contrary effects of RhoA and RhoC on barrier function. We hypothesized that each member of this subfamily can regulate these motor proteins in different regions of the human endothelial cell and then analyzed the effect of single Rho depletion on actomyosin by confocal microscopy. The lack of RhoC sharply reduced F-actin levels and preferentially decreased peripheral actin fibers (Fig. 6d, e). RhoB knockdown, which increases RhoA and RhoC expression levels but had no effect on barrier function, unexpectedly induced an overall increase in F-actin, compensated probably by a decrease of p-MLC, whereas RhoA silencing did not promote a significant actomyosin reorganization (Fig. 6d, e). Interestingly, the immunostaining of phosphorylated MLC in homeostatic conditions was preferentially localized at the periphery of the cell, from where it relocated to central stress fibers mostly upon RhoC knockdown, but not when RhoA and RhoB were depleted (Fig. 6d). In agreement with a RhoC-dependant regulation of peripheral actomyosin, only RhoC cells (Fig. 6f) and cells exposed to CN03 (Fig. 6g) exhibited robust perijunctional F-actin, confirming that modulating RhoC expression or Rho activity with CN03, promotes changes in the spatiotemporal localization of F-actin polymerization.

**Figure 6.**
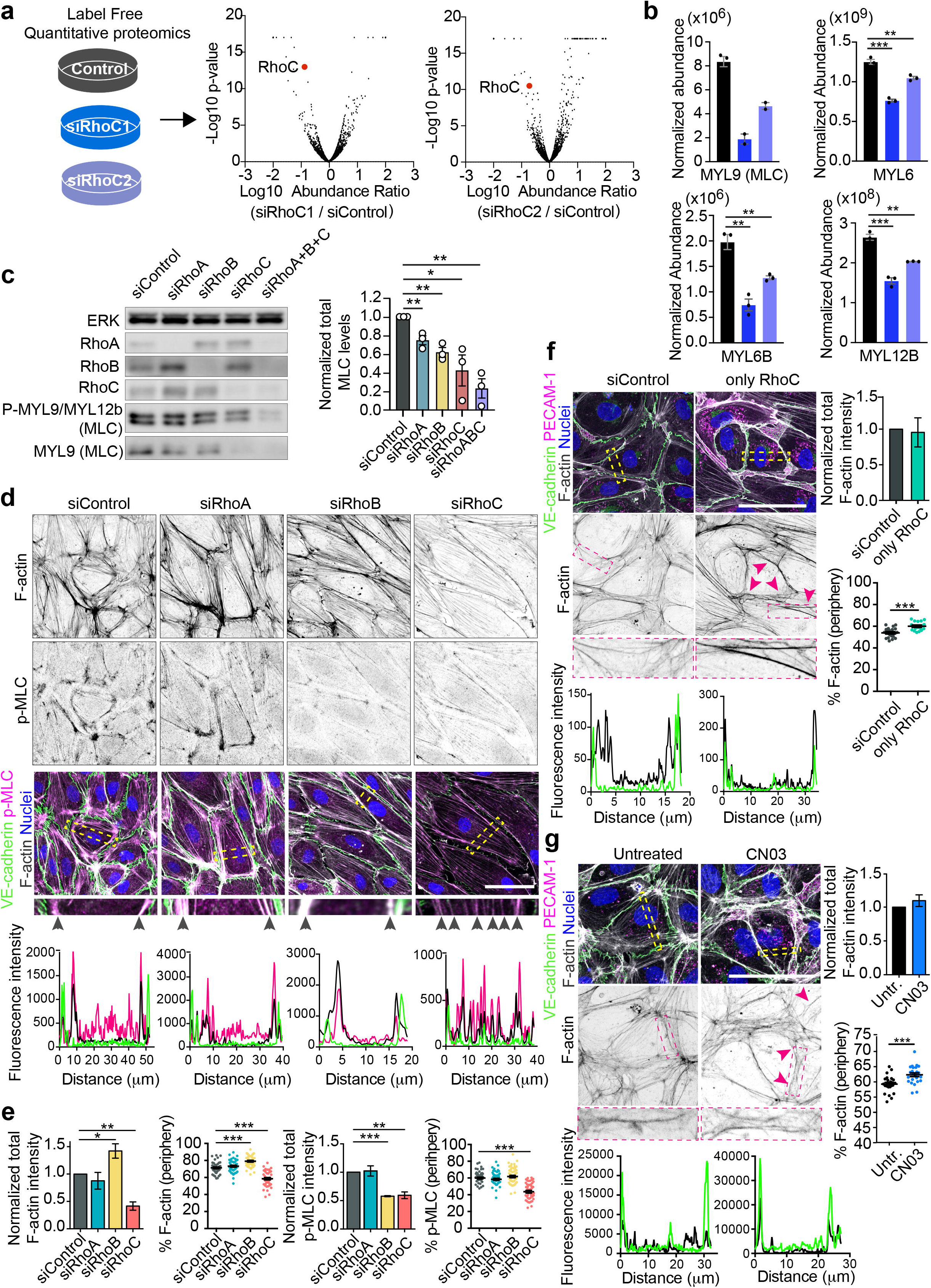
RhoC specifically regulates perijunctional actomyosin. **a,** Label-free quantitative proteomics of HUVECs transfected with the indicated siRNAs for 72 h. Volcano plots show the reduction of RhoC detected by quantitative proteomics as experimental control. **b,** Normalized abundance of protein components of actomyosin complexes obtained in the proteomic analysis. See Extended Data Table I. **c,** Reduction of MLC (MYL9) and pMLC (detecting both phosphorylated MYL9 and MYL12b) proteins expression in HUVECs transfected with the indicated siRNAs for 72 h. Right graph shows quantification of the relative levels of MLC with respect to siControl cells. **d,** Distribution of F-actin and p-MLC in confluent HUVECs transfected with the indicated siRNAs. Bottom graphs show the intensity profiles for VE-cadherin (green), p-MLC (magenta) and F-actin (black) corresponding to the squared areas in the images. Scale bar, 50 μm. **e,** Quantification of total and peripheral F-actin and p-MLC intensities. **f,** F-actin and junctional distribution in control and only RhoC HUVECs. Arrowhead point at junctional actin bundles. Bottom images show perijunctional F-actin details in a three-fold enlargement of the red boxed areas. Bottom graphs show the intensity profiles for VE-cadherin (green) and F-actin (black) corresponding to the yellow squared areas in the top images. Right graphs quantify total and perijunctional F-actin levels. Scale bar, 50 μm. **g,** F-actin and junctional distribution in Control and CN03-treated cells. Arrowhead point at junctional actin bundles. Bottom images show perijunctional F-actin details in a three-fold enlargement of the red boxed areas. Bottom graphs show the intensity profiles for VE-cadherin (green) and F-actin (black) corresponding to the yellow squared areas in the top images. Right graphs quantify total and perijunctional F-actin levels. Nuclei were stained with DAPI. Scale bar, 50 μm. Plots and graphs show the mean□±□SEM from at least three independent experiments. *, P< 0.05; **, P< 0.01; *** P<0.001.

It has been shown that MLC activation and phosphorylation can be mediated by the Rho kinase (ROCK), which phosphorylates MLC at the cell center, and the myosin light-chain kinase (MLCK), which localizes in cortical actin bundles and regulates MLC phosphorylation at cell borders ^42^. To test whether myosin-II activation and VE-cadherin expression levels are mechanistically coupled, we analyzed the effect of the ROCK inhibitor Y27632 and the MLCK inhibitor ML-7 on MLC phosphorylation, VE-cadherin levels and barrier function. Y27632 clearly reduced MLC phosphorylation, but had no major effect on VE-cadherin levels, consistent with findings in previous analyses ^15^. In contrast, the effect of the MLCK inhibitor ML-7 on MLC phosphorylation was not as strong as that of Y27632, but ML-7 clearly decreased VE-cadherin protein expression and provoked endothelial barrier dysfunction (Extended Data Fig. 7a, b). Accordingly, ML-7, but not Y27632, generated intercellular gaps (Extended Data Fig. 7c, arrowheads). Together, these experiments indicate that MLCK and the RhoC-mediated control of peripheral actomyosin regulate VE-cadherin expression and endothelial barrier function.

### RhoC is the only RhoA subfamily member that is homeostatically active and localized at cell borders

We next investigated why CN03 and RhoC strengthen the actin cytoskeleton close to cell-cell junctions. In addition, we also wondered why this recombinant protein has a net barrier-enforcing effect despite it similarly activates anti-barrier RhoA and RhoB and pro-barrier RhoC. We and other researchers had previously shown that unstimulated primary human endothelial cells express equal levels of RhoA and RhoC mRNA ^43^ and protein ^10^ but reduced levels of RhoB, whose expression is only increased upon long-term cytokine stimulation ^10^. We quantified the relative levels of activation of these three proteins by analyzing the ratio between the GTP-loaded Rho proteins, isolated by pull-down assays, and Rho protein expression detected in the initial cell lysates. We found that RhoC is almost eight times more active than RhoA and four times more active than RhoB (Fig. 7a). Since RhoB expression levels are very low relative to those of RhoA and RhoC in quiescent endothelial cells ^10^, these results show that RhoC accounts for most of the active RhoA subfamily proteins in unstimulated confluent human endothelial cells. Rho GTPases are translocated to the plasma membrane upon activation ^44^. In accordance to the previous result, RhoC, but not RhoA or RhoB, was found to be associated with a plasma membrane fraction in surface protein biotinylation assays, which enabled the fractionation of proteins exposed to the extracellular milieu along with proteins associated with them (Fig. 7b). This indicates that RhoC is the only one of these three GTPases that is mostly active under homeostatic conditions. Furthermore, while active, GTP-bound Rho GTPases are typically translocated to the plasma membrane, their GDP-bound, inactive forms complex with RhoGDI proteins in the cytoplasm, masking the Rho prenylated hypervariable region responsible for membrane translocation^44^. Immunoprecipitation assays revealed that RhoA was more strongly associated with RhoGDI compared to RhoC, consistent with the prevalent activation of RhoC relative to RhoA in confluent human endothelial barriers (Fig. 7c). It is of note that, RhoC also coprecipitated with RhoA, as recently described in the proximity interaction network of Rho GTPases ^45^, but not inversely. Finally, similar to the analyses previously performed in primary human vascular endothelial cells ^10^ and human corneal endothelial cells ^46^, we quantified the relative expression levels of endogenous RhoA, RhoB, and RhoC proteins in murine microvascular endothelial cells isolated from lungs. We found that RhoC is predominantly expressed compared to RhoA and RhoB, suggesting that this GTPase is central to microvascular endothelial cell function in murine experimental models as well (Fig. 7d). Collectively, these results demonstrate the dominant activation of RhoC with respect to its closest GTPases, particularly RhoA, which remains mostly inactive in homeostatic conditions. Since CN03 similarly activates RhoA, RhoB, and RhoC in human endothelial cells (Fig. 1a), the absolute number of activated RhoC molecules is far greater than those of activated RhoA and RhoB, tipping the balance of CN03 effect toward strengthening the endothelial barrier.

**Figure 7.**
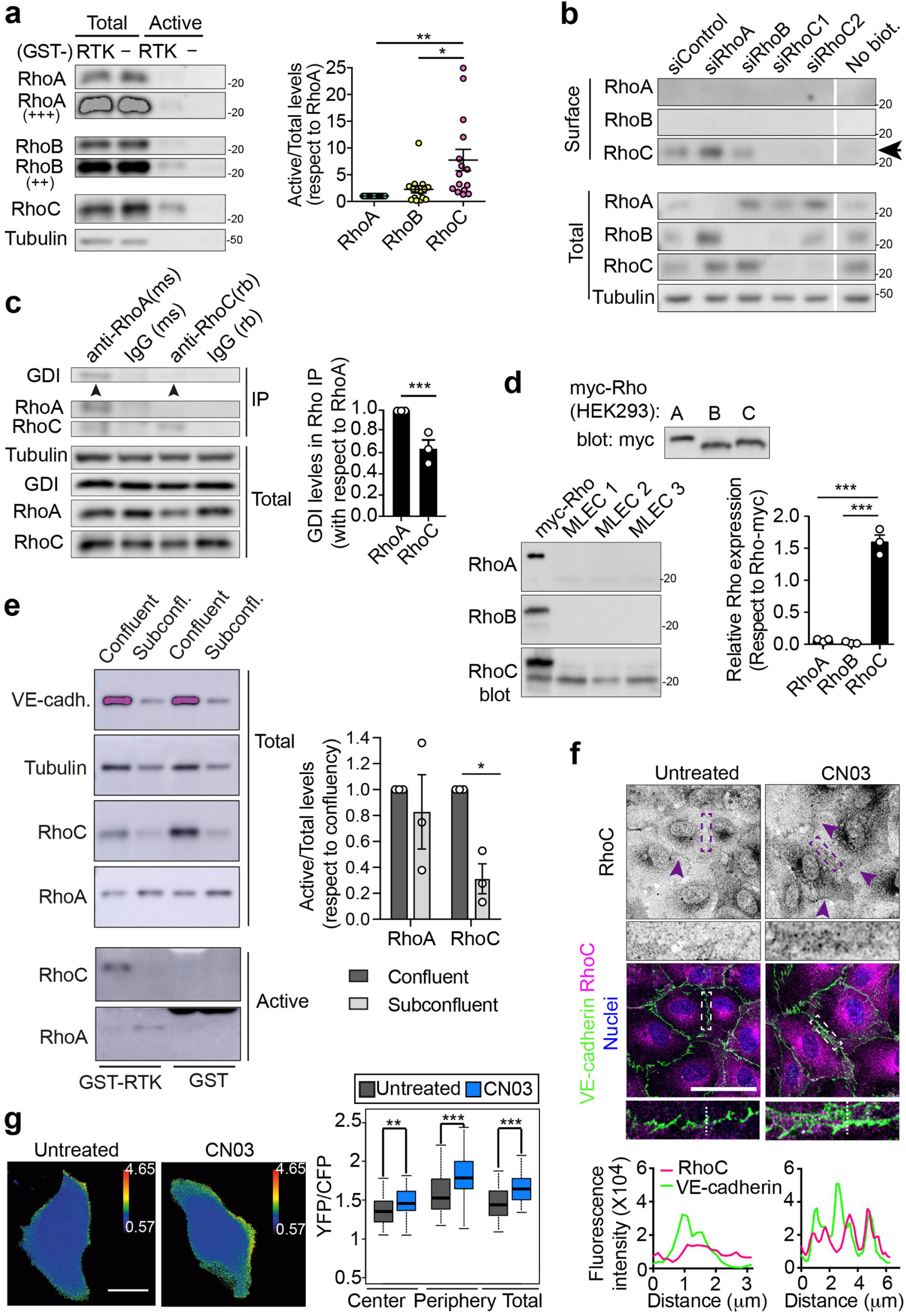
RhoC is the most active GTPase of the RhoA subfamily and is preferentially active at cell-cell junctions in confluent human endothelial cells. **a,** Relative activation levels of RhoA, RhoB y RhoC in unstimulated confluent HUVECs. Tubulin is shown as loading control. Right graph shows the quantification of Rho activation with respect to Rho expression normalized to the values obtained for RhoA. **b,** RhoC, but not RhoA and RhoB, is associated to surface plasma membrane proteins isolated by NeutrAvidin pull-down assays of cells previously biotinylated at the cell surface. **c,** Association of RhoA and RhoC to RhoGDI. Confluent HUVECs were lysed and subjected to immunoprecipitation with antibodies specific for RhoA and RhoC. Tubulin is shown as loading control. Bottom plots quantifies the relative levels of GDI association to RhoA and RhoC in the immunoprecipitates. **d,** Relative expression levels of RhoA, RhoB, and RhoC in mouse lung endothelial cells (MLEC). Three different lysates of HEK293 cells containing equal levels of exogenous RhoA, RhoB, or RhoC tagged with myc were generated as internal standards (top). Lysates of MLEC from three mice, together with the HEK293 lysate containing the corresponding exogenous myc-Rho (right), were blotted with antibodies against RhoA, RhoB, or RhoC. The expression level of each endogenous Rho protein in the MLEC lysates was then normalized to the intensity signal of their corresponding standard lysate and the loading controls, so the relative abundance of RhoA, RhoB and RhoC could be quantified (right graph). **e,** HUVECs were cultured at subconfluency or confluency for 72 h, lysed and subjected to pull-down assays with GST or GST conjugated to the RBD of rhotekin to detect active RhoA and RhoC. VE-cadherin is shown as a positive control of confluency-regulated protein expression. Tubulin is shown as a loading control. **f,** HUVECs were cultured at confluency for 72 h, exposed or not to CN03 for 3h, fixed and stained for RhoC (top images, inverted color) and VE-cadherin. Nuclei were stained with DAPI. Bottom images show a three-fold enlargement of the squared areas. Bottom graphs show the intensity profiles of the discontinuous lines in the enlarged images. Scale bar, 50 μm**. g,** FRET analysis of RhoC-FLARE in the presence or absence of 1.5 μg/ml CN03. Bottom images show quantification of the FRET signal increase in different cellular regions. Plots show the mean□±□SEM from at least three independent experiments. *, P< 0.05; **, P< 0.01; *** P<0.001.

Interestingly, cell confluency significantly increased the expression and activation of RhoC, but not of RhoA, suggesting that homeostatically active RhoC is regulated by contacts between endothelial cells (Fig. 7e). Indeed, confocal analyses with specific antibodies revealed that RhoC was evenly distributed throughout the cell, but becoming more concentrated at cell borders in response to CN03 (Fig. 7f, enlarged area). Junctional RhoC was also observed in only RhoC cells, whereas RhoA staining was not confined to any cellular structure in any cell type. RhoB distribution was mostly in endosomes in siControl and only RhoB cells (Extended Data Fig. 8a), as previously described ^10^. We further examined the effect of CN03 on the localization of RhoC activity by carrying out a FRET analysis with the RhoC FLARE biosensor, which consists of the Rho binding domain of ROCK (RBD) (residues 905-1046), conjugated with full-length RhoC and flanked by monomeric fluorescent proteins able to undergo FRET when RBD binds activated RhoC (Extended Data Fig. 8b) ^47^. As a prior control, we observed higher FRET in RhoC constitutively active (Q63L) FLARE and a lower FRET in RhoC dominant negative (T19N) FLARE, than in RhoC FLARE (Extended Data Fig. 8c). FRET between the two fluorophores was preferentially detected close to the border of endothelial cells expressing RhoC FLARE (Fig. 7g). Moreover, CN03 selectively increased FRET signal at the cell periphery, consistent with the translocation observed for endogenous RhoC, and indicating that this chimeric protein promotes RhoC activation at cell borders. Thus, RhoC is the most active GTPase in confluent endothelial monolayers, its expression is regulated by cell-cell junctions and is preferentially active at cell borders.

### The junctional localization and high activity of RhoC depend on the R188 residue from the hypervariable region

Collectively, or results show that the activity of RhoC is significantly higher than that of RhoA and RhoB, probably due to the concentration of various Rho GEFs at cell-cell junctions, where RhoC concentrates ^48–50^. We investigated the molecular basis of the contrasting differences between RhoC and RhoA and hypothesized that these differences lie in their C-terminal hypervariable regions, as these are their most divergent protein domains and contain the prenylation site that enables lipid modifications and localization to the plasma membrane upon activation. RhoA and RhoC have polybasic sequences adjacent to their prenylation C residue, but RhoA also contains a phosphorylatable S residue that is absent from RhoC ^51^. First, we conjugated these hypervariable regions to the C-terminus of GFP (GFP-RhoA-hyper and GFP-RhoC-hyper) to determine whether they are sufficient to target this fluorescent protein to VE-cadherin-positive membrane domains (Fig. 8a). GFP-RhoC-hyper partially colocalized with VE-cadherin to a greater extent than GFP-RhoA-hyper. As expected, when the prenylation C-residue was mutated to G (GFP-RhoA/C-hyper^C190G^), the two chimeric GFP constructs became mostly cytoplasmic. Substituting the R188 residue of RhoC by an S, as occurs in the hypervariable region of RhoA (GFP-RhoC-hyper^R188S^), reduced the junctional localization of the fluorescent chimera. Reciprocally, substituting the S188 residue of the RhoA segment by an R, as in the RhoC hypervariable domain, (GFP-RhoA-hyper^S188R^), increased junctional staining, indicating that this position is fundamental for Rho localization. Mutating these residues in the full GFP-RhoC proteins had a similar effect in the junctional localization of the protein (Fig. 8b) This outcome was not specific to the S substitution, since GFP-RhoC^R188A^ displayed a reduction in junctional localization with respect to GFP-RhoC similar to that of GFP-RhoC^R188S^, suggesting that the positive charge of R188 residue is fundamental to targeting RhoC to the membrane. Importantly, the alterations in membrane localization of these mutants were associated with changes in RhoC activity, with those that had less junctional localization being less active (Fig. 8c). Moreover, overexpression of GFP-RhoC, but not of GFP-RhoA or GFP-RhoC^R188S^, reinforced endothelial barrier function (Fig. 8d and Extended Data Fig. 9a), revealing the pivotal role of R188 in the specific pro-barrier effect of RhoC. It is of note that the reciprocal mutation in GFP-RhoA, GFP-RhoA^S188R^, increased its junctional localization but did not increase its activity indicating that other molecular features intervene in the regulation of this GTPase (Extended Data Fig. 9b, c). Thus, the R188 residue contributes to the functional opposition between RhoC and its counterparts in the RhoA subfamily, which can be exploited to strengthen the vascular endothelial barrier.

**Figure 8.**
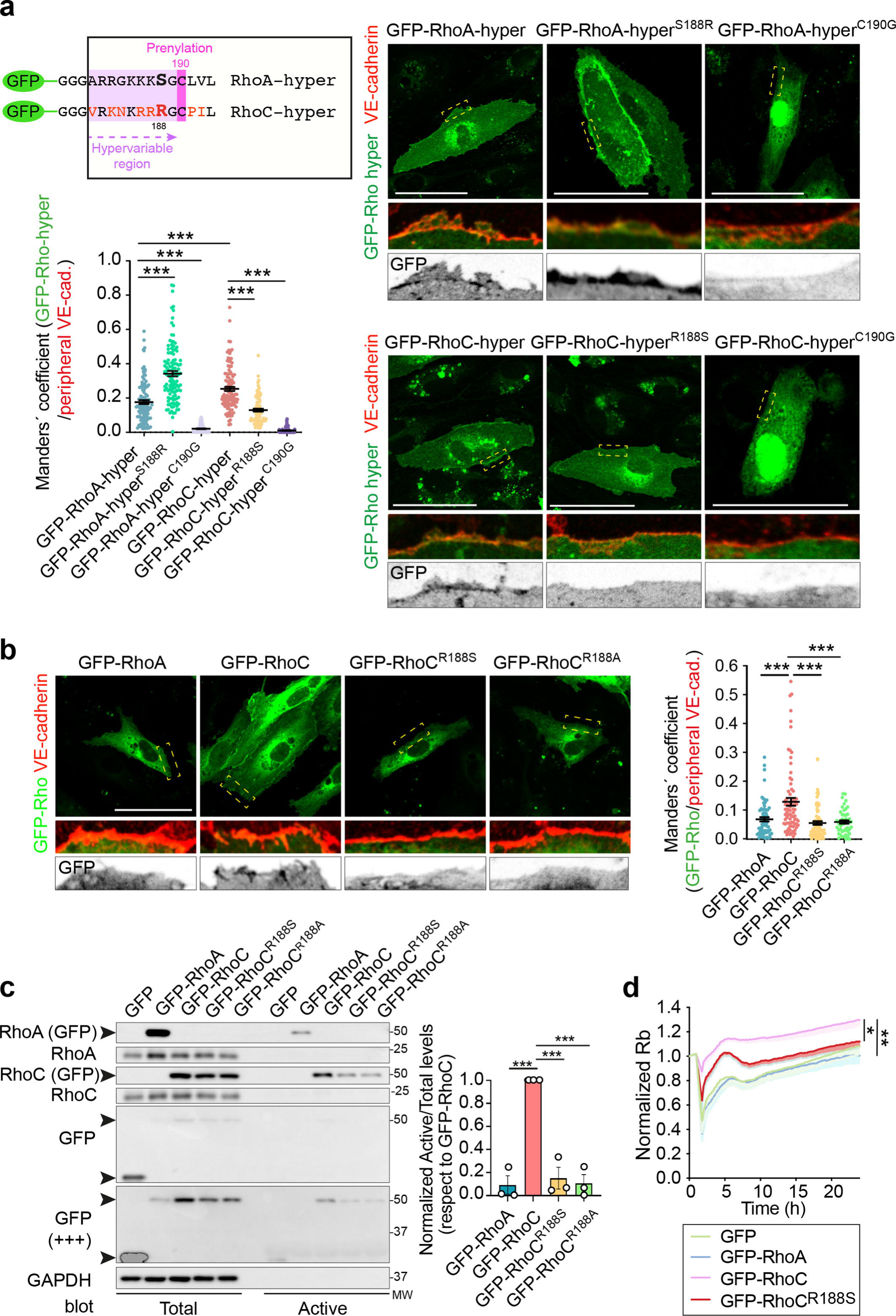
The junctional localization and the pro-barrier role of RhoC are regulated by a R188 residue in the hypervariable region, which is not present in RhoA. **a,** Chimeric constructs of GFP conjugated to the RhoA and RhoC hypervariable regions (GFP-Rho-hyper). RhoA contains a S residue two positions upstream the prenylation site, whereas RhoC contains a R residue. GFP-RhoA-hyper and GFP-RhoC-hyper containing the wild type hypervariable regions, as well as mutated sequences in which the RhoA S residue was substituted by R (GFP-RhoA-hyper^S188R^) and the RhoC R residue replaced by S (GFP-RhoC-hyper^R188S^) were expressed in HUVECs for 24 h, cells fixed and stained for VE-cadherin. Bottom left plots show the Manders’ coefficients for GFP-hyper chimeras and VE-cadherin. **b,** The RhoC R residue was substituted by S or the uncharged amino acid A in GFP-RhoC constructs. Expression plasmids containing the indicated chimeras were ectopically expressed for 24 h, cells were fixed, stained for VE-cadherin and junctional co-localization quantified with Manders’ coefficient. **c,** HUVECs expressing the indicated GFP-Rho proteins were lysed and subjected to pull-down assays with GST-RTK to detect GTP-loaded RhoA and RhoC. Total and active fractions were immunoblotted with the indicated antibodies. Right graph shows the relative quantification of each GFP-Rho activation with respect to GFP-RhoC values. **d,** HUVECs transiently expressing the indicated GFP-Rho proteins were plated on ECIS arrays and TEER and Rb analyzed. Plots show the mean□±□SEM from at least three independent experiments. *, P< 0.05; **, P< 0.01; *** P<0.001.

## DISCUSSION

Our findings reveal a surprising divergence in function among the highly similar Rho GTPases, RhoA, RhoB, and RhoC. RhoC exhibits unique regulatory and functional features in human endothelial cells: it is regulated by cell confluency, localizes to and stabilizes cell-cell junctions and junctional actomyosin, and is much more active in homeostatic confluent endothelial cells than the closely related RhoA and RhoB. Importantly, the predominant role of this GTPase represents a novel therapeutic target since it can be exploited to reinforce the endothelial barrier and protect the vasculature, opening promising new avenues for developing treatments against acute pathologies linked to endothelial dysfunction.

In the canonical model of regulation of endothelial barrier function by Rho GTPases, the RhoA subfamily drives contraction and disruption of barrier integrity, whereas Rac subfamily members, namely Rac1, are involved in barrier reformation and maintenance. Disruptive stimuli such as thrombin, LPS and histamine activate RhoA ^6,7,10^ whereas pro-barrier stimuli such as sphingosine-1-phospate and hepatic growth factor activate Rac^8,9,22,52^ . However, a dual role in permeability, promoting both pro-barrier and anti-barrier signaling, has already been identified in epithelial cells for the RhoA subfamily, yielding important mechanistic insights ^53–55^. Sahai and Marshall have shown that ROCK is the main effector mediating RhoA-mediated acute contraction and barrier disruption, whereas mDia is the effector involved in the RhoA subfamily-mediated stabilization of epithelial AJs ^56^. In our experimental settings we have also found that ROCK is involved in acute cell contraction ^10,15^ but, as we show here, is not involved in RhoC-dependent maintenance of endothelial barrier integrity. Instead, MLCK appears to be mediating this latter effect, implying that, in these vascular cells, the signaling pathways also diverge downstream the RhoA subfamily.

A RhoA subfamily pro-barrier function already holds significant pathological relevance in intestinal epithelia: RhoA inactivation is a major pathogenic factor that causes loss of barrier integrity in inflammatory bowel disease ^57^. The RhoA signal is impaired in intestinal epithelial cells (IECs) from patients with inflammatory bowel disease (IBD), which causes progressive epithelial barrier dysfunction. Furthermore, conditional loss of RhoA was sufficient to mimic chronic intestinal inflammation, impair intestinal barrier function and provoke an increase in IEC shedding. Although the specific effects of RhoA KO on epithelial cell-cell junctions, cytoskeletal rearrangement and IES permeability were not addressed, treatment with CN03 clearly reduced intestinal inflammation and the IBD score in a genetic model altering RhoA activity ^57^.

Septicemia causes a paradigmatic endothelial barrier disruption that leads to vascular leakage, organ edema and failure, making endothelial cells an important therapeutic target ^1,58^. However, the progress in finding new treatments focused on the host response have been at standstill for the last decade for this pathology. Current strategies of hemodynamic stabilization that directly target the microvasculature are limited to the use of vasopressors, general anti-inflammatory drugs such as corticosteroids that may regulate vascular beds, and other alternative treatments to reduce the inflammatory mediators in the bloodstream ^59–61^. It is noteworthy that some attempts to reinforce vascular barrier function targeting endothelial molecular scaffolds have been carried out. Imatinib, an inhibitor of Abl kinases, broadly used for cancer treatments, has been found to activate Rac1 and strengthen endothelial cell adhesion to the extracellular matrix and hence, the endothelial barrier function, in preclinical models of sepsis ^62^. Indeed, imatinib has been tested in some clinical trials with moderate positive effects on ameliorating inflammatory systemic diseases such as severe COVID19 ^63,64^. We have compared the effects of imatinib and CN03 on vascular leakage *in vivo* (not shown) and found a similar reduction of hyperpermeability between these two compounds, which suggests that RhoA subfamily of GTPases controls a signaling pathway that could also be exploited as a new therapeutic option to prevent organ damage upon systemic inflammation. Whether RhoC-mediated and Rac1-mediated pathways crosstalk at any point under these circumstances is an open question that needs to be addressed, although we have experimentally addressed the effect of RhoA, RhoB and RhoC silencing on Rac1 activity and found no significant changes. This implies that imatinib and CN03 control parallel signaling routes, opening the possibility of targeting simultaneously both of them and investigate their additive effects.

The early reports on the specific effect of RhoA subfamily member overexpression present somewhat contradictory results. RhoC is more disruptive for cell-cell junctions than exogenous RhoA expression in the human colon epithelial cell line HCT116 ^56^, whereas RhoC expression reinforces the perijunctional ring in the human colon epithelial cell line T84, suggesting a pro-barrier role for this GTPase ^55^. Most of these studies have been performed overexpressing mutants, which increased transversal stress fibers and induced cell contraction, instead of externally modulating endogenous RhoA activity. This suggests that activity turnover and subcellular localization are essential for endogenous Rho proteins to function correctly. In this respect, and also in the epithelium, the outcome of Rho GTPase activation is known to be strongly determined by its subcellular location. In particular, local, perijunctional RhoA activation supported by junctional myosin-II improves barrier function in epithelia. Such activation close to cell-cell junctions could be prevented by myosin IIa knockdown and myosin inhibition with blebbistatin, which disrupts AJs in a similar way to that we reported for endothelial cells with the MLCK inhibitor ML-7 ^53^. Putting these and our results together, an appealing concept emerges with respect to epithelial and endothelial junctions as mesoscopic hubs that orchestrates differential modulation of Rho signaling by regulating the accessibility of Rho regulators. Junctional protein complexes recruit RhoGEFs ^65–69^ and RhoGAPs ^70^, but they could also prevent the accessibility of these regulators, as it happens for the Rho activity suppressor p190B RhoGAP in columnar epithelial cells ^53^. These accessibility is regulated, for example, by mechanotransduction ^68^ and may be a relevant target for endothelial hyperpermeability caused by pathological inflammation ^50,71,72^

Considering the importance of endothelial junctions as Rho regulatory mesoscopic domains, we have identified the molecular basis of RhoC junctional localization, which underlies the functional differences between RhoA and RhoC in these cells. The positively charged R188 in RhoC is a S188 in RhoA, which can be phosphorylated by PKA and thus negatively charged ^51^. R188 significantly contributes to target RhoC to cell-cell junctions, thereby allowing this GTPase to gain access to junctional domain enriched in Rho GEFs and perhaps devoid of Rho GAPs. Indeed, the formation of cell-cell junctions upon confluency increases the activity and expression of RhoC, but not of RhoA. This major role of confluency in shifting the balance of expression and activity RhoA and RhoC can have potential consequences in processes involving junctional remodeling such as angiogenesis and vasculogenesis, which should be thoroughly explored in the future. However, in the mature microvasculature, our findings have unveiled that RhoC is the main homeostatic regulator of barrier function and a potential target for therapies that ameliorate endothelial barrier damage and edema during systemic inflammation.

## METHODS

### KEY RESOURCES TABLE

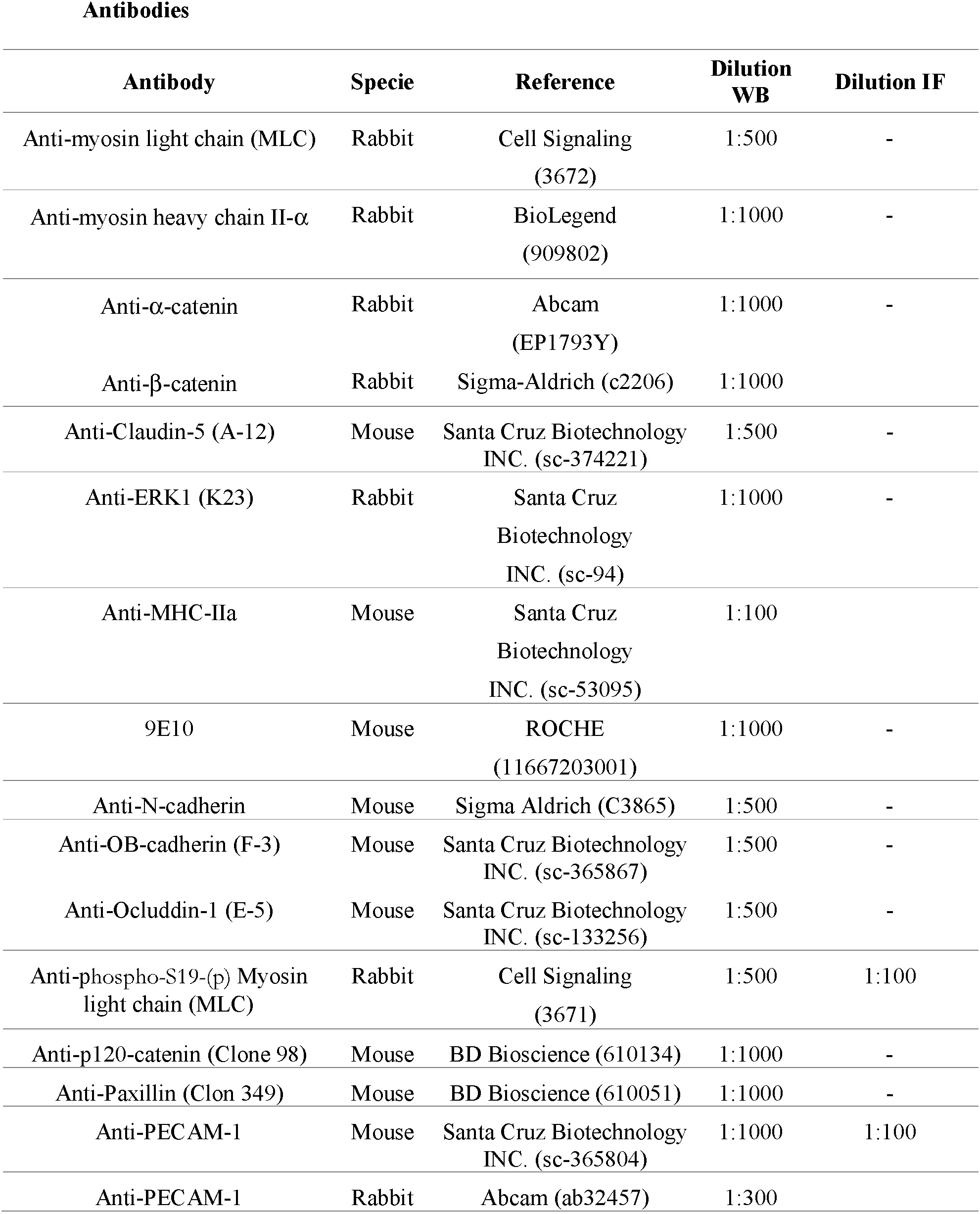
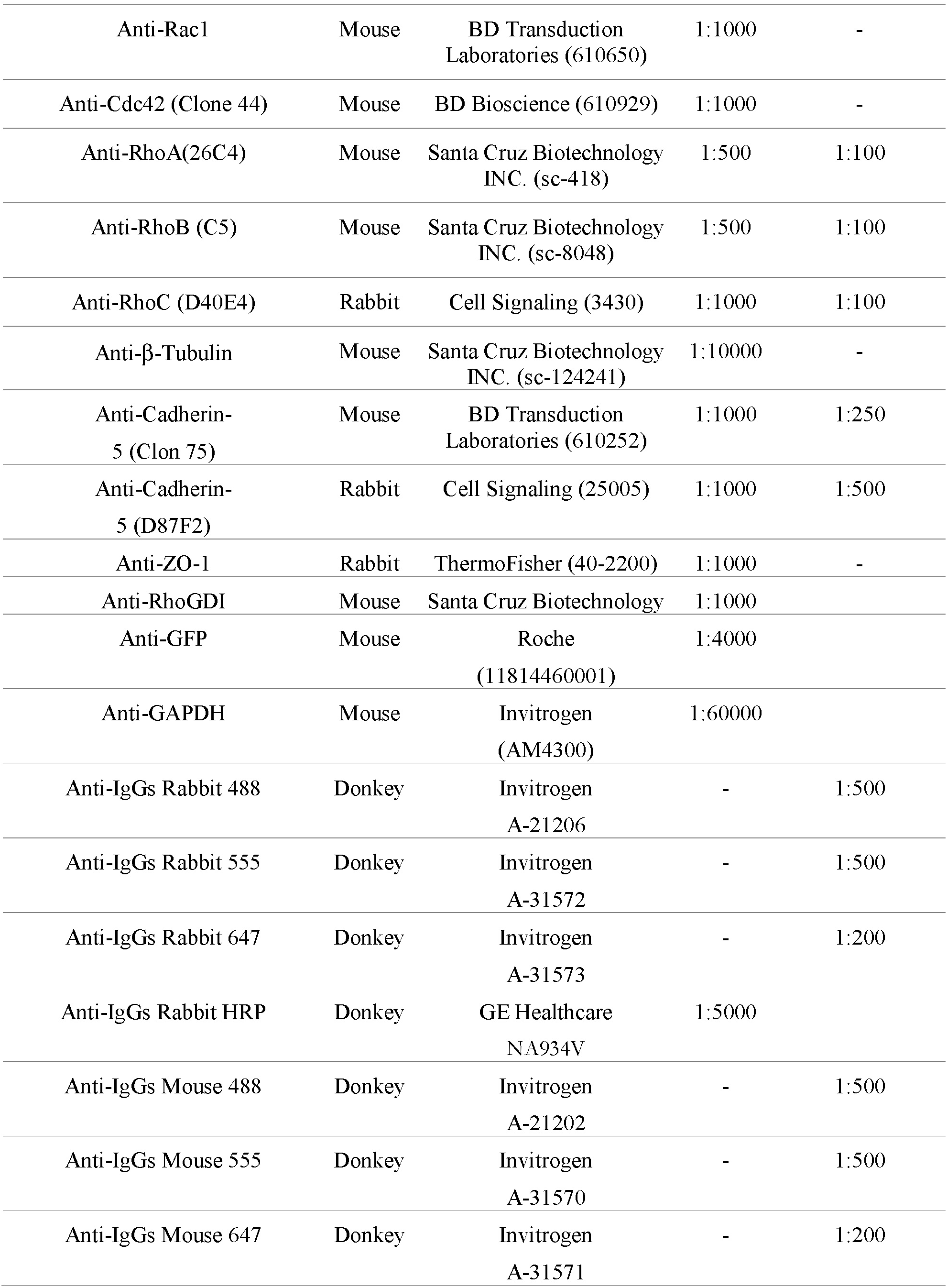
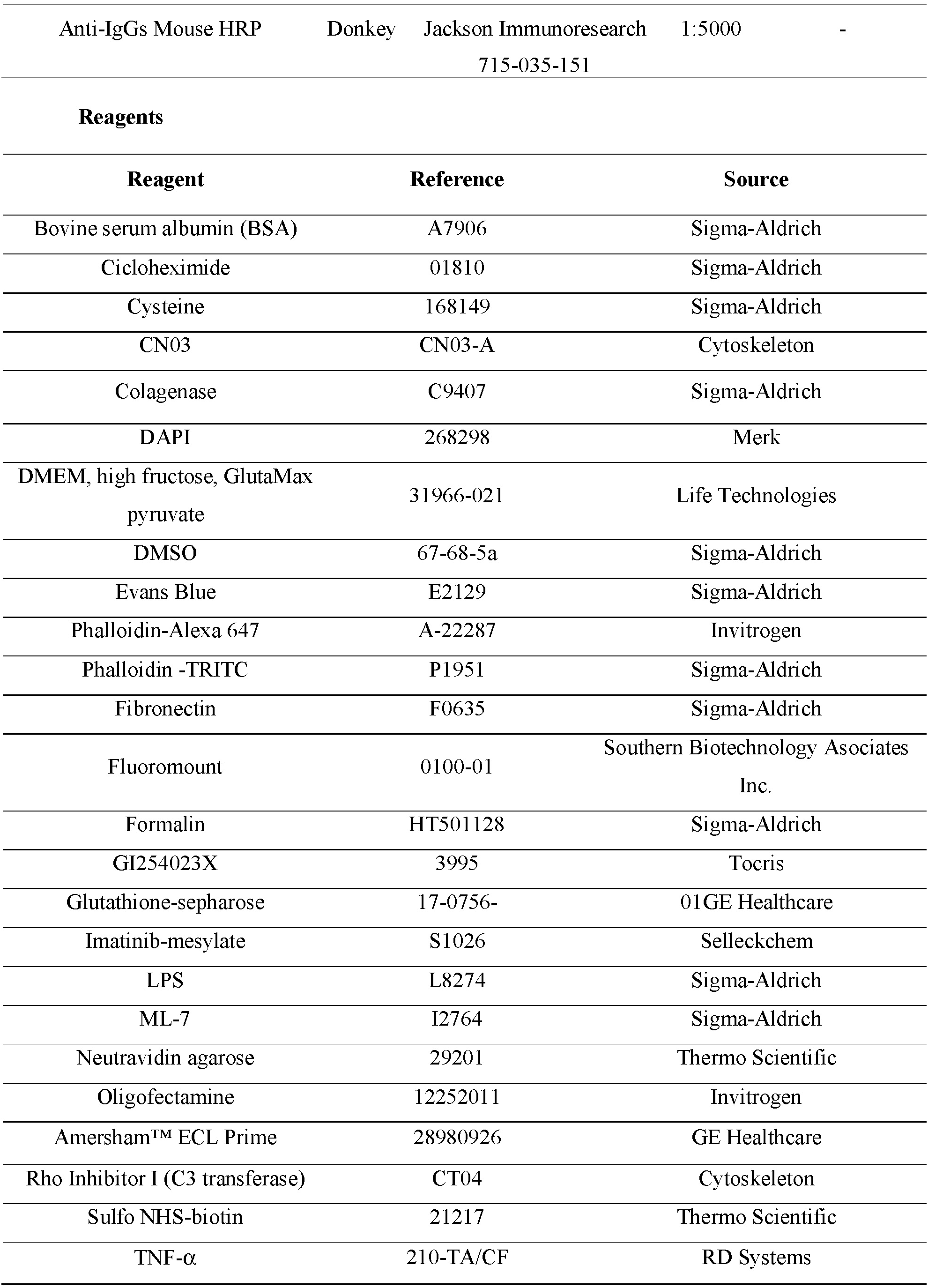
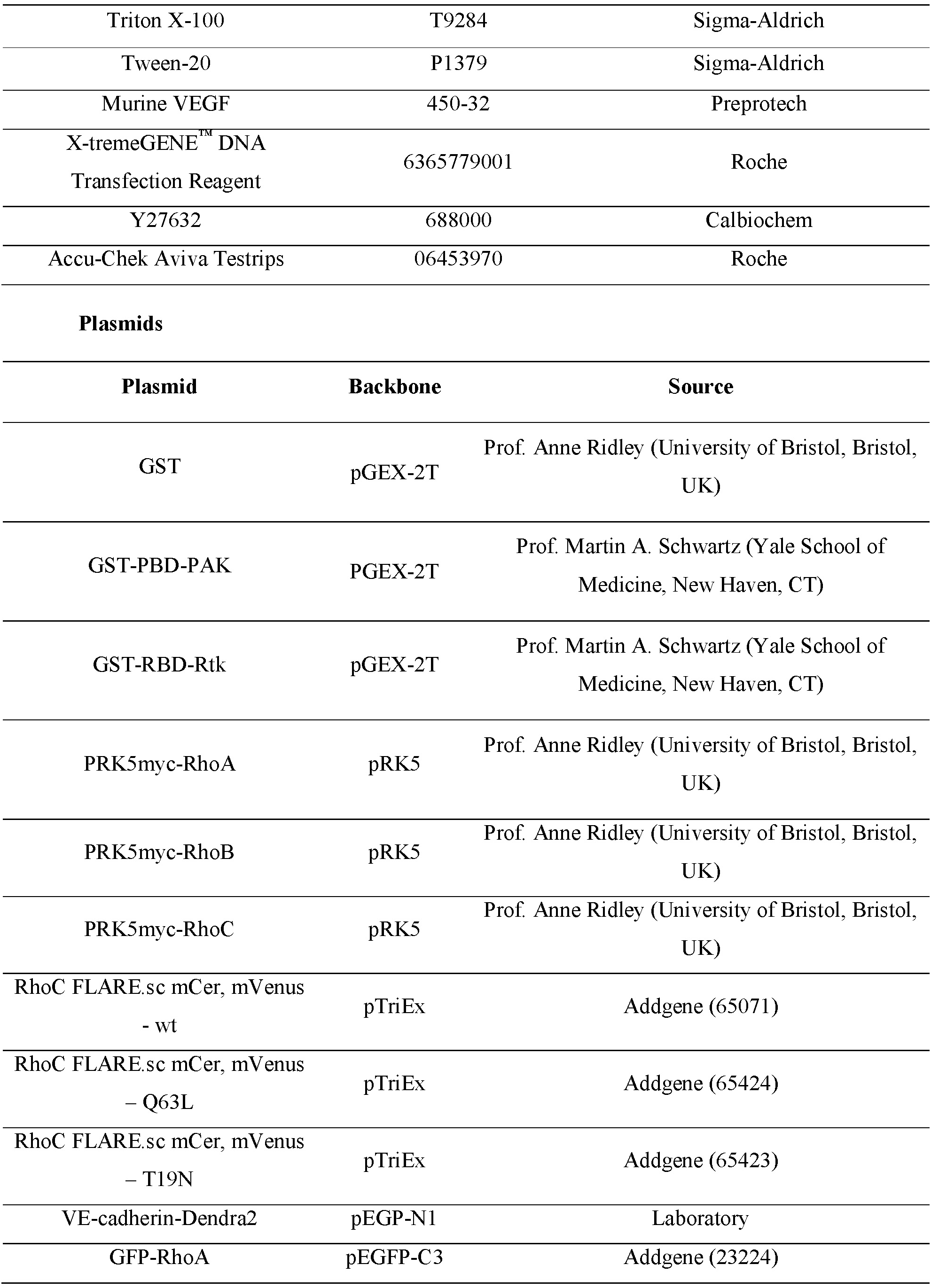
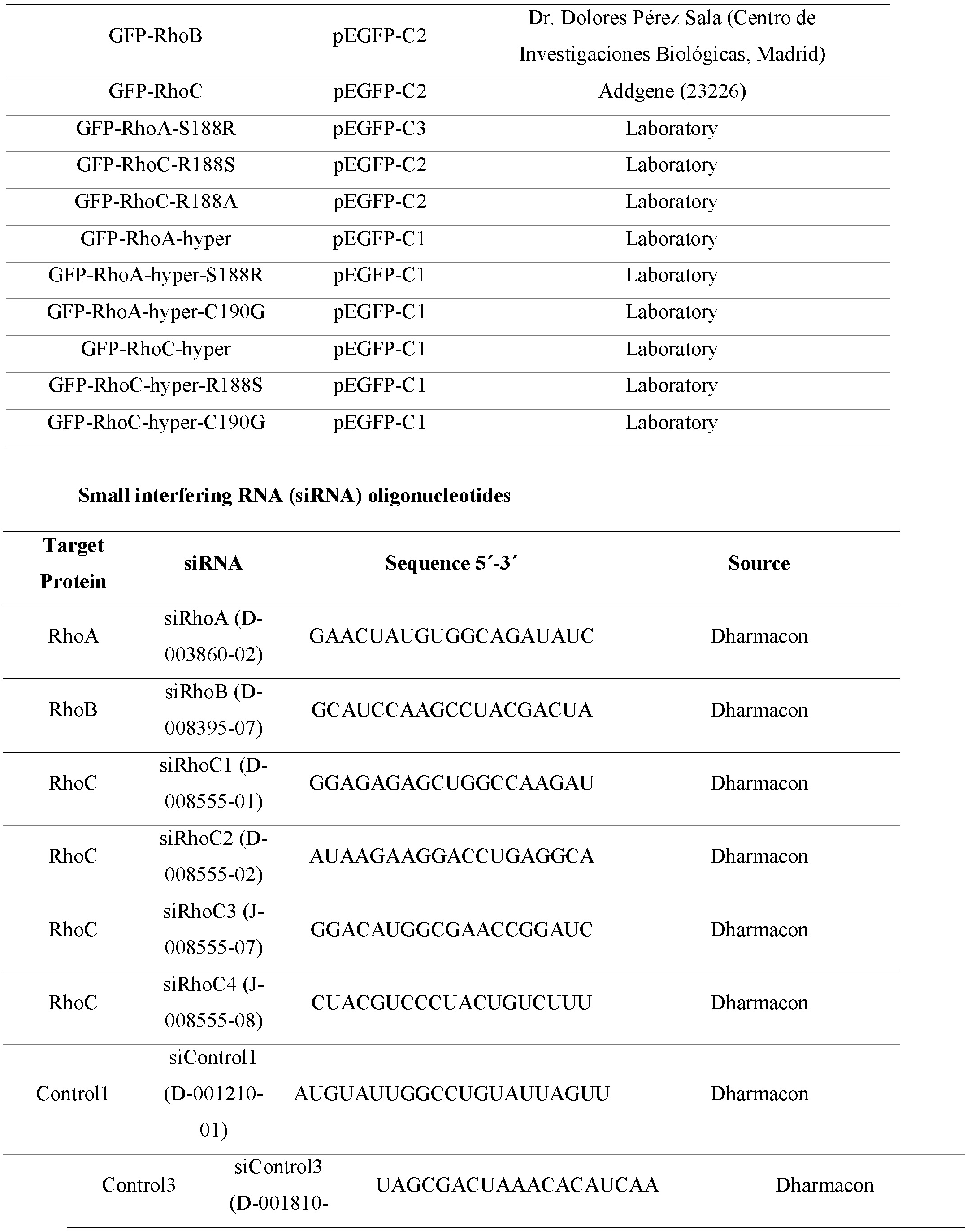
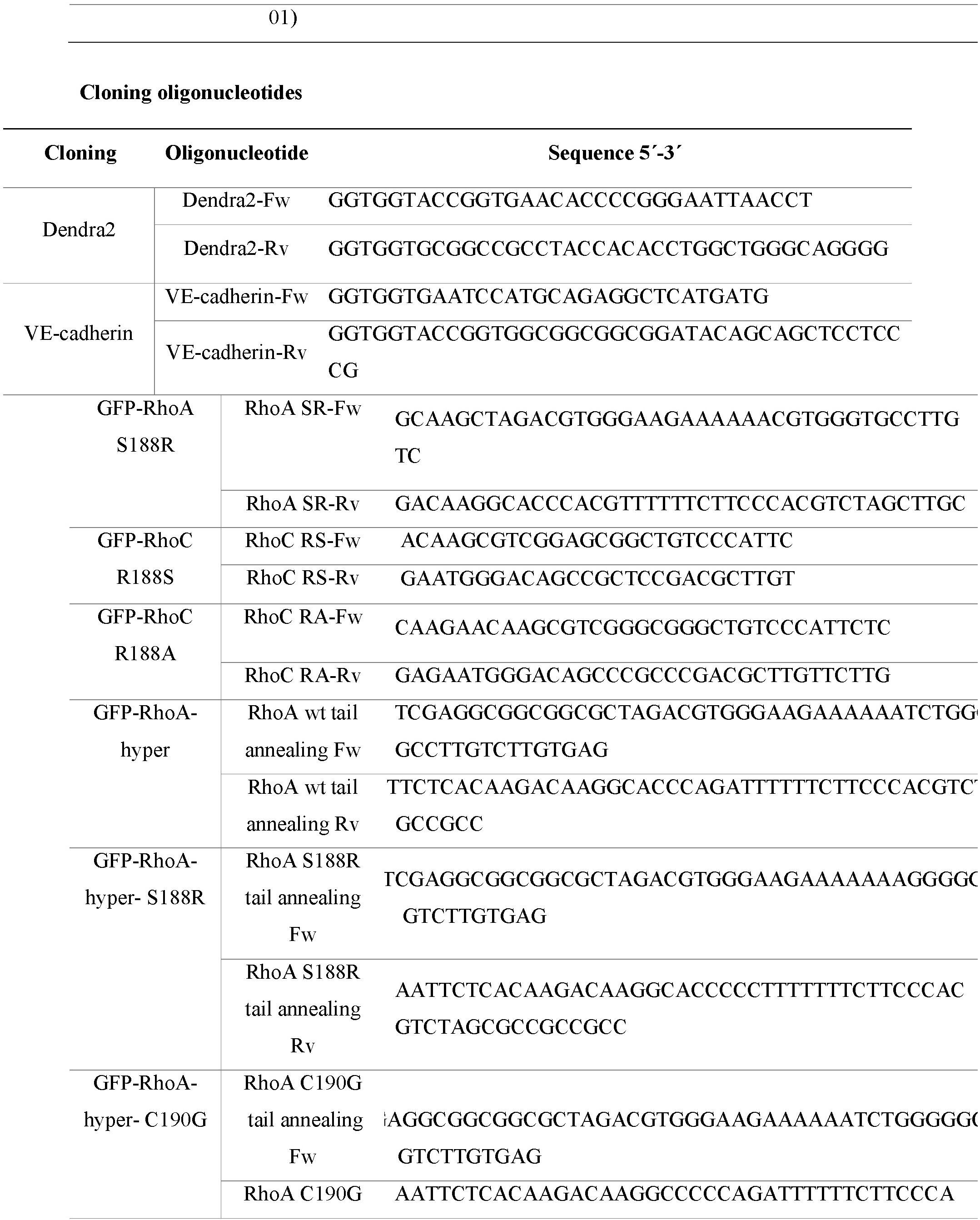
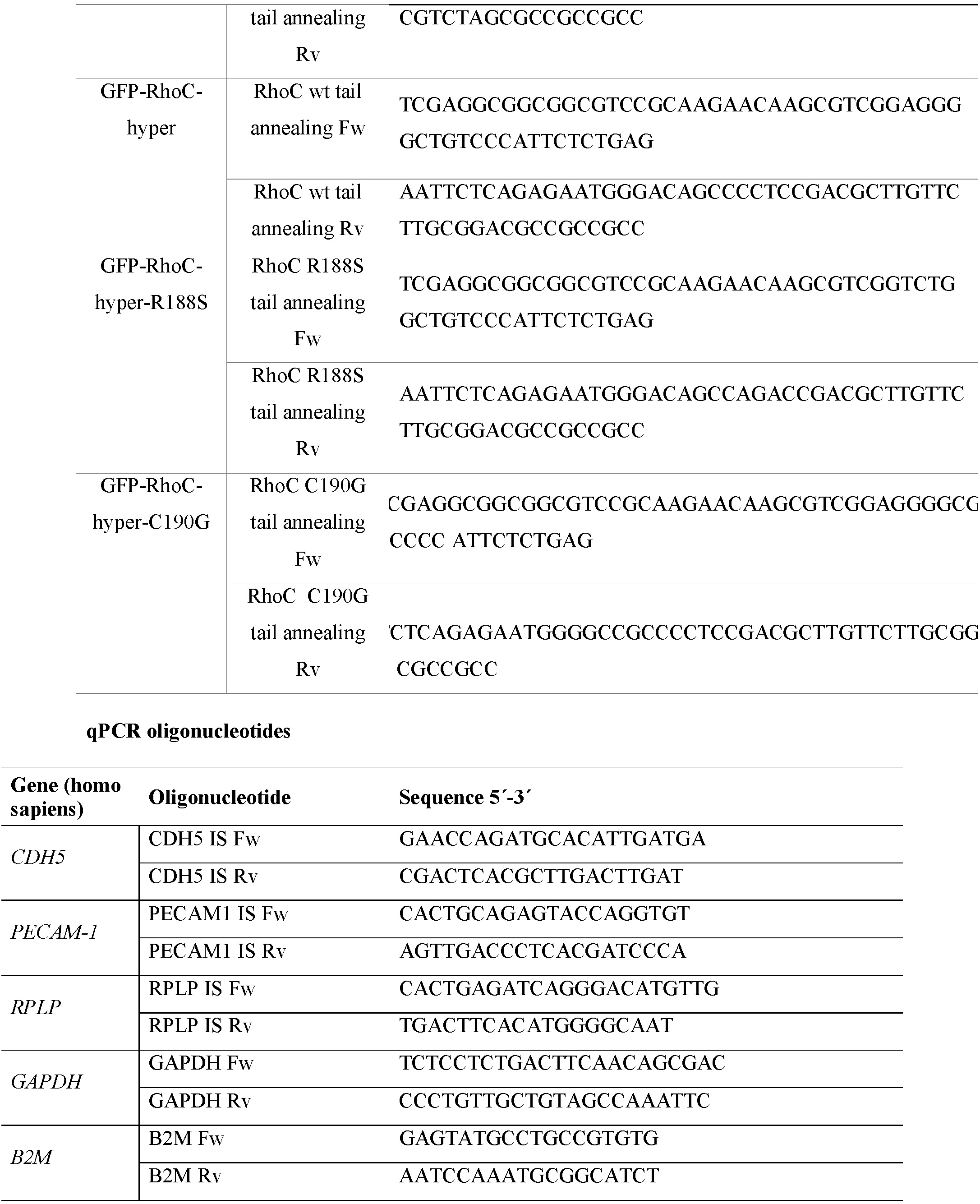

#### Cell culture and transfection

Human embryonic kidney (HEK) 293 cells were obtained from the American Type Culture Collection and grown in DMEM supplemented with 10% FBS. HUVECs were purchased from Lonza and grown in EGM-2 medium (Lonza). HDMVECs and the corresponding culture medium (CADMEC medium) were purchased from TebuBio and grown according to the manufacturer’s instructions. EA.hy.926 cells were grown in DMEM supplemented with 10% FBS in an atmosphere of 5% CO2/95% air. EA.hy.926 cells were a gift from C.J.S. Edgell (University of North Carolina, Chapel Hill, NC). Plates were precoated with 10 μg/ml fibronectin (Sigma-Aldrich) for at least 30 min (plastic) or 2 h (glass coverslips) prior to cell culture. All cell types were grown at 37°C in an atmosphere of 5% CO2/95% air. Murine lung endothelial cells were isolated as previously described ^21^. Briefly, mouse lungs were minced and digested with 0.1% type I collagenase solution (Gibco) for 1 h at 37 °C. Lung tissue was further disaggregated with a 19.5 G needle and subsequently filtered through a cell strainer. Lung cell suspension was then subjected to positive sorting with an antibody against the endothelial cell marker ICAM-2 conjugated to magnetic Dynabeads Sheep Anti-Rat IgG (11035) from Thermo Fisher, for 30 min at 4 °C. After extensive washes, endothelial cell rich fraction was lysed with Laemmli sample buffer and subjected to western blot analysis.

For siRNA transfection, HUVEC or HDMVECs were plated at subconfluence (5×10^5^ cells per 60 mm-dish) in their respective culture media without antibiotics. Cells were transfected 24 h later with a mixture of 8 μl of Oligofectamine (Invitrogen) and the indicated siRNA to a final concentration of 100 nM. 24 h after oligotransfection, cells were trypsinized and plated at confluence in different wells for parallel assays, such as western blotting, immunofluorescence or permeability assays, performed 72 h after transfection.

For DNA transfection, 2×10^5^ cells per well were seeded on 24-well plates containing fibronectin-coated glass coverslips. For time-lapse analysis of living cells, including FRET and photoconversion assays, 2×10^5^ HUVECs were seeded on fibronectin-coated µ-Slide 8 Well chambers (Ibidi). Cells were transfected with a mixture of 1 μg of plasmid DNA and 1 μl X-tremeGENE™ HP DNA Transfection Reagent (Sigma-Aldrich) per well. Medium was replaced 24 h after transfection and experiments were performed 48 h post-transfection.

#### Molecular cloning of GFP-hyper constructs

RhoA and RhoC hypervariable regions were added to the C-terminal domain of GFP in a GFP-C1 vector using the using the annealed oligonucleotides listed in the Key Resources Table. The inserted sequences contain one Xho-I site at the 5’end one EcoR1 site at the 3’ end, which allowed the ligation of the annealed oligonucleotides to the GFP plasmid.

#### Site-directed mutagenesis of RhoA and RhoC hypervariable regions

The aforementioned mutations in the hypervariable regions of RhoA-GFP and RhoB-GFP were generated according to the manufacturing instructions of the QuickChange Lightning Site-Directed Mutagenesis kit (Agilent Technologies) using the mutagenic primers listed in the Key Resources Table. Plasmid containing the indicated mutations were transformed into XL10-Gold ultracompetent cells. Plasmids from selected colonies were sequenced before scaling up their purification with EndoFree Plasmid Maxi kit (Qiagen) and subsequent transfection.

#### qRT-PCR

Oligonucleotides for qPCR are listed in the Key Resource Table. 1 μg RNA from HUVECs was subjected to reverse transcription with the High-Capacity RNA-cDNA kit (Applied Biosystems). RT-qPCR was performed from the resulting cDNA in a CFX 384 thermocycler (Bio-Rad) using the SsoFast EvaGreen Supermix (Bio-Rad) and forward and reverse primers, designed with Probefinder software (Roche). Parallel qPCR assays for the housekeeping genes RPLP and GAPDH and B2M were performed to normalize data from each point. Three replicates of all the samples were run in parallel with non-reverse transcription controls for all the targets, which yielded no detectable amplification. Unless otherwise indicated, the quantification cycle (Cq) values fell in the 20–24 range, which denotes medium to high levels of these transcripts. Results were quantified and represented (2−ΔCq) using GenEx software, correcting for the efficiency of each pair of primers. The analyses were performed in technical triplicates. Contamination with genomic DNA was negligible. Subsequently, these values were normalized with respect to the mean values of housekeeping genes used as reference. The stability of the candidate reference genes was probed with geNorm and Normfinder algorithms within GenEx, and the combination of the two most stable genes was chosen against which to normalize the results.

#### Endothelial barrier function assay

Trans-endothelial electric resistance (TEER) assays with an electric cell-substrate impedance sensing system (ECIS 1600R; Applied Biophysics) were performed as described ^4,10,21^. TEER indicate the resistance values at 4000 Hz. The Rb, α, and membrane capacitance values (Cm) were modelled using the ECIS software (Applied Biophysics) running the experiment in the multi-frequency mode, which ranges between 250 and 64000 Hz. This mathematical modelling split TEER values into α values, which correlates with focal adhesion formation, Rb values, which reflects paracellular barrier formation (Rb), and Cm, which reflects the membrane capacitance TEER decrease was quantified as the percentage of decrease in the TEER, taking as 100% the difference between the TEER value at t = 0 h and the TEER value of an empty well.

#### Western blot

For Western blot analysis, equal amounts of proteins were loaded onto polyacrylamide gels (8-12%) under reducing conditions and transferred to Immobilon-P membranes (EMD Millipore). After blocking with 5% BSA (wt/vol) and 0.05% (vol/vol) Tween-20 in TBS (25 mM Tris, pH 7.4, and 150 mM NaCl), membranes were incubated overnight with the indicated antibodies, washed with TBS containing 0.05% Tween 20, and incubated for 30 min with the corresponding secondary antibodies coupled to HRP. After extensive washes, the signal was visualized with an enhanced chemiluminescense Western blotting kit (ECL; GE Healthcare). Band intensities were quantified using Fiji and the results were represented relative to controls.

#### Surface biotinylation assays

HUVECs were plated in 60-mm dishes and incubated with 0.250 mg/ml sulfo-NHS-biotin in PBS supplemented with 1.5 mM Ca^2+^ and Mg^2+^ (PBS-Ca^2+^ Mg^2+^) for 30 min at 4°C. Cells were washed with PBS-Ca^2+^ Mg^2+^to eliminate the excess of non-labelled NHS-biotin and incubated for 10 min with DMEM 10% FBS to quench the remaining free NHS-biotin, and then washed again with PBS-Ca^2+^ Mg^2+^. Cells were lysed with were harvested in lysis buffer (1x TBS, 10 mM EDTA, 1% Triton, 0.3 μg/ml Caliculin A, 1 mM PMSF, 10 μg/ml Aprotinin, Leupeptin and Pepstatin, 1 mM Sodium Orthovanadate, 10 mM β-Glycerol Phosphate, and 10 mM Sodium Fluoride) and passed through a 0.6-mm 24G needle. All of these procedures were performed at at 4°C. Subsequently, lysates were incubated 5 min at 37°C and centrifuged at 14,000 rpm for 5 min. Part of the supernatant was conserved as total lysate and the remainder was subjected first to a preclearing step with agarose for 1 h at 4°C and then to pull-down assay with NeutrAvidin agarose (Thermo Scientific) for 2 h at 4°C to isolate biotinylated proteins.

#### Activation of endogenous RhoA subfamily of GTPases and pull-down assay for measuring GTP-loaded Rho proteins

HUVECs were grown at confluency for 3 days and starved for 24 h. Cells were incubated with 1.5 μg/ml CN03 for 3 h and then lysed and subjected to pull-down assays to measure GTP-loaded RhoA, RhoB and RhoC, as previously described ^10,73^ . HDMVECs were stimulated with 5 μg/ml CN03 after previous titration assays. Briefly, cell lysates were incubated at 4 °C in assay buffer with 10 μg of glutathione-sepharose beads conjugated with GST-RBD-Rtk to isolate active RhoA, RhoB and RhoC, or conjugated to GST-PBD-PAK to isolate active Rac1 and Cdc42. A fraction of the initial postnuclear supernatants and the final GST pull-down samples were subjected to SDS-PAGE and western blot analysis with the indicated anti-Rho antibodies.

#### Laser scanning confocal microscopy

Confluent cells grown on coverslips were fixed in 4% paraformaldehyde (PFA) for 15 min, and rinsed and treated with 10 mM glycine for 2 min to quench the aldehyde groups. Cells were then permeabilized with 0.2% Triton X-100, and rinsed and blocked with 3% bovine serum albumin (BSA) in PBS for 15 min at room temperature (RT). Cells were incubated for 30 min with the primary antibodies, rinsed in PBS, and incubated for 30 min with the appropriate fluorescent secondary antibodies. Actin filaments were detected with fluorophore-conjugated phalloidin (see key resources table). Incubation with antibodies and other fluorescence reagents was always carried out at 37°C. Phosphorylated proteins were stained using Tris-buffered saline (20 mM Tris, pH 7.4; 150 mM NaCl) instead of PBS. Confocal laser scanning microscopy was carried out with a confocal Zeiss LSM710 system coupled to an AxioImager M2 microscope, a confocal Zeiss LSM 800 system coupled to an AxioObserver microscope, and a confocal Nikon AR1+ system coupled to an Eclipse Ti-E microscope. Fiji was used for quantification of the area covered by AJs. Using Z-stacks of endothelial monolayers projected in a single image, a Region Of Interest (ROI) was drawn to encompass the entire cell including the intercellular junctions. This provides us with the total intensity of the protein of interest in the cell as well as the cell area. A smaller adapted ROI was drawn to surround the cell inside beneath the intercellular junctions. Values from the inner area were subtracted from those from the outer, peripheral area, providing a measurement of the percentage of overlapping reticular areas with respect to the cellular area, as well as the junctional distribution of the indicated proteins with respect to the total cellular staining. The average intensity was measured in each region by dividing the RawIntDen value (sum of the pixels in a given region) by the area, allowing for comparison of parameters between different images.

#### Generation of VE-cadherin-Dendra2 construct, photoconversion and time-lapse microscopy

VE-cadherin-GFP-N1 vector previously generated in the laboratory from a VE-cadherin expression plasmid kindly donated by Elisabetta Dejana. Photoconvertible VE-cadherin-Dendra2 chimera was generated by substituting the GFP sequence by the Dendra2 sequence. Briefly, Dendra2 was amplified by PCR using the oligonucleotides described in the Key Resources Table, containing the Age-1 and Not-1 restriction sites. To study VE-cadherin dynamics, HUVECs were transfected with the expression vector coding for VE-cadherin-Dendra2 in µ-Slide 8 Well (Ibidi). 48 h later and 5 h before photoconversion, the medium was replaced with EBM-2 without phenol red, supplemented with 1% FBS and 10 mM Hepes, pH 7.4. Time-lapse microscopy was performed with a confocal Nikon AR1+ system coupled to an Eclipse Ti-E microscope, in an environmental chamber at 37°C and an atmosphere of 95% O2/5% CO2. Cells were simultaneously filmed in green (λ = 488 nm) and red (λ = 543 nm). An image was acquired before photoconversion as the initial fluorescence value for both wavelengths. A region of interest (ROI) in cell-cell junctions was then irradiated with a 405-nm laser at 10% power. The ROI underwent an emission shift from 488-nm to 543-nm. Images were acquired every 2 s for 8 min. To analyze the VE-cadherin-Dendra2 experiments, we quantified the mean fluorescence intensity in the photoconverted area over time using Fiji. We extracted disappearance rate constants from exponential curves of VE-cadherin-Dendra2 543-nm intensity decay.

#### FRET imaging of RhoC activity

The human endothelial cells Ea.hy.926 were plated in a µ-Slide 8 Well chamber (Ibidi) and transfected with the corresponding RhoC-FLARE. 24 h post-transfection, cells were starved for 24 h in EBM-2 without phenol red, supplemented with 1% FBS and 10 mM HEPES. CN03 was added 3 hours prior to the experiment. Images were acquired in an environmental chamber at 37°C and an atmosphere of 95% O2/5% CO2, using a wide-arc lamp fluorescence microscope (FRET system). Emission ratio imaging was performed with a 460-500-nm excitation laser/520-550-nm emission YFP. We previously defined the range between the active and inactive states of the biosensor by analyzing FRET efficiency in cells expressing the active and dominant negative forms of RhoC-FLARE. Images were processed and analyzed using Fiji software.

#### Transmission electron microscopy and 3D image reconstruction

Confluent human lung microvascular endothelial cells transiently expressing PECAM-1-GFP were in situ fixed with 4% paraformaldehyde and 2% glutaraldehyde in 0.1 M phosphate buffer, pH 7.4 for 2 h at RT. Postfixation was carried out with a mixture of 1% osmium tetroxide and 0.08% potassium ferricyanide for 1 h at 4°C and then with 2% uranyl acetate for 1 h at RT. Samples were dehydrated with ethanol and processed for standard Epon (TAAB-812) embedding. After polymerization, consecutive orthogonal ultrathin sections (70 nm) were collected on formvar-coated slot grids and stained with uranyl acetate and lead citrate. Finally, sections were examined in a Jeol JEM-1400Flash transmission electron microscope operating at 100 kV. Images were recorded with a Gatan OneView (4K x 4K) CMOS camera at 4000x magnification.

For 3D reconstruction, electron microscopy images were binned 4 times, with an effective pixel size at the specimen level of 11.2 nm, and their intensities were normalized to the same mean and standard deviation. The electron microscopy images of the series were mutually aligned with the IMOD software tool ^4^. To this end, structural features unequivocally identified in adjacent sections, especially membrane invaginations and multivesicular bodies, were manually selected (around 25-30 per each pair of images) and employed as fiducial marks to align each pair of consecutive sections independently. Finally, each section was transformed into a common alignment in which the section located in the middle of the series acted as a reference, resulting in the aligned series and thus yielding the 3D reconstruction. Modelling of the 3D reconstruction was done with IMOD by manually tracing the plasma membranes. The total thickness of the 3D reconstruction consisting of 27 individual sections was around 1.89 μm.

#### Intradermal injection of inflammatory stimuli and Miles assay

Female CD-1 mice were intraperitoneally injected with saline or 0.175 mg/kg of CN03. A Miles assay was performed after 3.5 h after injection. Briefly, we depilated the back of the mice and retro-orbitally injected them with 150 μl 0.5% w/v Evans Blue (Merck). 15 min later, saline, 20 μg of LPS and 50 ng of VEGF were intra-dermally injected in the back of each mouse for 15 min. Mice were then euthanized and the stimulated points of the skin, as well as 100 mg of liver, lung and kidney were harvested from each mouse. Evans blue was extracted with formamide at 55 °C for 48 h, measured spectrophotometrically and normalized with respect to the tissue weight.

#### LPS-induced systemic inflammation

Female CD-1 mice were intraperitoneally injected with saline as control or 50 μg/g LPS to mimic the final state of sepsis disease in combination or not with 0.175 mg/kg or 0.525 mg/kg (X3) of CN03. A Miles assay was performed 17 h later in the indicated organs as described, which revealed an increase of tissue edema in response to LPS injection. Blood samples were collected before and after the treatment for each animal. TNF and IL-6 were measured by flow cytometry and plasma glucose was measured with Accu-Chek Aviva Testrips (Roche).

#### Mass spectrometry analysis of CN03 sequence

CN03 was analyzed by LC-ESI-MSMS mass spectrometry at the Proteomics Service of the National Center for Biotechnology (CSIC, Spain). 1 μg CN03 was run on a 12% denaturing polyacrylamide SDS-PAGE gel. A single gel band was detected and cut, deposited in a 96-well plate, and automatically processed in a Proteineer DP (Bruker Daltonics, Bremen, Germany). The digestion protocol used was described by Shevchenko et al. with the following minor variations: gel plugs were first washed with 50 mM ammonium bicarbonate and then with acetonitrile before reduction with 10 mM DTT in 25 mM ammonium bicarbonate; alkylation was performed with 55 mM IAA in 50 mM ammonium bicarbonate. Then, the gel pieces were washed with 50 mM ammonium bicarbonate and 100% acetonitrile, and then dried under a nitrogen stream. Modified porcine trypsin (sequencing grade; Promega, Madison WI) was added at a final concentration of 16 ng/μl or 25 ng/μl, respectively, in 25% acetonitrile, 50 mM ammonium bicarbonate, and digestion was carried out for 6 hours at 37°C. The reaction was stopped by adding 0.5% TFA for peptide extraction. The tryptic peptides were dried by vacuum centrifugation and resuspended in mobile phase A (0.1% formic acid in water) for LC ESI-MSMS. The NanoLC ESI-MSMS analysis of tryptic peptides was performed using an Eksigent 1D Plus nanoLC coupled to a SCIEX 5600 TripleTOF QTOF mass spectrometer. The analytical column used was a reverse-phase C18 silica-based column of 75 µm x 15 cm, particle size of 1.7 µm, and pore size of 100 Å (Waters). The trap column was a PepMap C18 (Dionex, Sunnyvale, California), particle diameter of 5 µm, pore size of 100 Å, connected online with the analytical column. The loading pump supplied a solution of 0.1% trifluoroacetic acid in 98% water/acetonitrile 2% (Merck) at 3 µL/min. The nanoflow pump provided a flow rate of 250 nL/min and operated under gradient elution conditions, using 0.1% formic acid (Fluka, Buchs, Switzerland) in water as mobile phase A and 0.1% formic acid in 100% acetonitrile as mobile phase B. The elution gradient was performed according to the following scheme: isocratic conditions of 96% A: 4% B for 5 min, linear increase to 40% B in 25 min, linear increase to 95% B in 1 min, isocratic conditions of 95% B for 5 min, and return to initial conditions in 10 min. The injection volume was 5 µl. The nanoLC system was coupled via a nano-spray source (SCIEX) to the QTOF mass spectrometer operating in positive ion mode with the capillary voltage set at 2600 V. Data-dependent acquisition allowed sequentially obtaining both full scan (m/z 350-1250) MS spectra followed by tandem MS CID spectra of the 25 most abundant ions. Dynamic exclusion was applied to prevent the isolation of the same m/z for 20 seconds after its fragmentation.

The MS and MS/MS data obtained were processed using PeakView (SCIEX). For protein identification, MSMS spectra were searched against a UniProtKB (http://www.uniprot.org) database consisting of entries from Proteobacteria (38594387 entries). Database searches were performed using a licensed version of Mascot v.2.4 (www.matrixscience.com; Matrix Science, London, UK). Search parameters were set as follows: carbamidomethyl cysteine as a fixed modification and oxidized methionine and acetyl (protein N-term) as variable modifications. The peptide mass tolerance was set to 25 ppm for MS1 and 0.05 Da for MS/MS spectra, respectively, and 1 missed cleavage was allowed.

#### Label-free quantitative proteomics

Tryptic digestion and tandem mass spectrometry analysis. Identical amounts of each replicate were individually digested with trypsin in reducing and alkylating (chloroacetamide) conditions. Tryptic peptides were dried in a speed-vacuum system and desalted using C18 tips. Thereafter, they were subjected to independent LC-MS/MS analysis using a nano liquid chromatography system (Ultimate 3000 nano HPLC system, Thermo Fisher Scientific) coupled to an Orbitrap Exploris 240 mass spectrometer (Thermo Fisher Scientific). Samples (5 μL) were injected on a C18 PepMap trap column (5 µm, 300 µm I.D. x 2 cm, Thermo Scientific) at 20 µL/min, in 0.1% formic acid in water, and the trap column was switched on-line to a C18 PepMap Easy-spray analytical column (2 µm, 100 Å, 75 µm I.D. x 50 cm, Thermo Scientific). Equilibration was done in mobile phase A (0.1% formic acid in water), and peptide elution was achieved in a 120 min gradient from 4% - 50% B (0.1% formic acid in 80% acetonitrile) at 250 nL/min. Data acquisition was performed in DDA (Data dependent acquisition) mode, in full scan positive mode. Survey MS1 scans were acquired at a resolution of 60,000 at m/z 200, scan range was 375-1200 m/z, with Normalized Automatic Gain Control (AGC) target of 300 % and a maximum injection time (IT) of 45 ms. The top 10 most intense ions from each MS1 scan were selected and fragmented by Higher-energy collisional dissociation (HCD) of 30%. Resolution for HCD spectra was set to 15,000 at m/z 200, with AGC target of 100 % and maximum ion injection time of 80 ms. Precursor ions with single, unassigned, or six and higher charge states from fragmentation selection were excluded.

Protein identification and label-free protein quantitation analysis. Raw files were processed using Proteome Discoverer (PD) version 2.5 (Thermo Fisher Scientific). MS2 spectra were searched using a combination of four search engines (Mascot v2.7, MSFragger, Amanda and Sequest), against a target/decoy database built from sequences corresponding to the *Homo sapiens* reference proteome downloaded from Uniprot Knowledge database. Database included typical laboratory contaminants. Search parameters considered fixed carbamidomethyl modification of cysteine, and the following variable modifications: methionine oxidation, possible pyroglutamic acid from glutamine at the peptide N-terminus and acetylation of the protein N-terminus. The peptide precursor mass tolerance was set to 10 ppm and MS/MS tolerance was 0.02 Da, allowing for up to two missed tryptic cleavage sites. Spectra recovered by a False Discovery Rate (FDR) <= 0.01 filter were selected and the signals corresponding to each identified peptide were extracted, normalized, assigned by protein inference to their corresponding proteins and used for quantitative analysis. Label-free Protein quantification used unique + razor peptides. The abundance parameter, which correlates with the sum of the intensities of the peptides identified for each protein was analyzed to establish significant changes in protein expression.

#### Patient Samples and Study Approval

The study was approved by the local Ethics Committee (La Paz University Hospital, Madrid, PI-4100 version 2, 2020). All the participants provided written consent and the data were treated according to recommended criteria of confidentiality, following the ethical guidelines of the 1975 Declaration of Helsinki. Patients >18 years who met the diagnostic criteria for sepsis according to the Third International Consensus Definitions for Sepsis and Septic-Shock ^75^ were enrolled in this study on arrival at Accident and Emergency Service of La Paz University Hospital. Healthy volunteers were recruited from the Blood Donor Services of La Paz University Hospital. Details of the blood collection, serum preparation and circulating levels of cytokines were described in detail elsewhere ^4^.

#### Statistical analysis

Data from at least three independent experiments are expressed as the mean and standard error of the mean (SEM). An unpaired Student’s t-test or two-way ANOVAs were used to establish the statistical significance (p < 0.05) of group differences between the means using Prism 7.0 software. Data were obtained from at least three independent experiments.

## AUTHOR CONTRIBUTIONS

N.C-A. and J.M. contributed to the conception and experimental design. N.C-A., P.M-P., M.G-F., A.C., C.C-N., S.B. and G.dR. performed the experiments and acquired the data. M.G-F carried out the RhoC-FRET FLARE analysis. N.C-A., A.C. and C.R. carried out all the *in vivo* experiments. N.C-A. and C.C-N. carried out the photoconversion assays. A.P. performed all the proteomic analyses. J.J.F. performed the 3D reconstruction of the EM images. N.C.A. and J.M. analyzed the data. E.L-C. M.F. and I.J-A. provided material support and reviewed the manuscript. J.M. wrote the manuscript.

## ACKNOWLEDGMENTS

The expert technical advices of the confocal microscopy, electron microscopy, genomic and animal facilities are gratefully acknowledged. The work was supported by grants PID2020-119881RB-I00 from AEI (to CC-N, PM-P, NC-A, SB, GdR, GC-T, and JM) and Grant TomoXliver2, Ref: S2022/BMD-7232 funded by Comunidad Autónoma de Madrid. (to SB and JM), and IND2019/BMD-17139 (to GC-T and JM) from Comunidad de Madrid. This research work was also funded by the European Commission─NextGenerationEU (Regulation EU 2020/2094), through CSIC’s Global Health Platform (PTI Salud Global) (to JM). SB is also supported by Endocornea2, Convenio Colaboración CSIC, funded by Instituto de Investigación Fundación Jiménez Díaz. The authors also thank Dr Phil Mason, who provided English language support. The authors declare that no conflicts of interest exist.

## REFERENCES

1. Goldenberg, N. M., Steinberg, B. E., Slutsky, A. S. & Lee, W. L. Broken barriers: a new take on sepsis pathogenesis. Sci Transl Med 3, 88ps25 (2011).

2. Fajgenbaum, D. C. & June, C. H. Cytokine Storm. N Engl J Med 383, 2255–2273 (2020).

3. Rittirsch, D., Flierl, M. A. & Ward, P. A. Harmful molecular mechanisms in sepsis. Nat Rev Immunol 8, 776–787 (2008).

4. Colás-Algora, N. et al. Simultaneous Targeting of IL-1–Signaling and IL-6–Trans-Signaling Preserves Human Pulmonary Endothelial Barrier Function During a Cytokine Storm. Arterioscler Thromb Vasc Biol 43(11), 2213–2222 (2023).

5. Teuwen, L.-A., Geldhof, V., Pasut, A. & Carmeliet, P. COVID-19: the vasculature unleashed. Nat Rev Immunol 20, 389–391 (2020).

6. Mikelis, C. M. et al. RhoA and ROCK mediate histamine-induced vascular leakage and anaphylactic shock. Nat Commun 6, 6725 (2015).

7. Amerongen, G. P. van N., Delft, S. van, Vermeer, M. A., Collard, J. G. & van Hinsbergh, V. W. M. Activation of RhoA by Thrombin in Endothelial Hyperpermeability. Circ Res 87, 335–340 (2000).

8. Garcia, J. G. N. et al. Sphingosine 1-phosphate promotes endothelial cell barrier integrity by Edg-dependent cytoskeletal rearrangement. J Clin Invest 108, 689–701 (2001).

9. Komarova, Y. A., Mehta, D. & Malik, A. B. Dual Regulation of Endothelial Junctional Permeability. Sci. STKE 2007, (2007).

10. Marcos-Ramiro, B. et al. RhoB controls endothelial barrier recovery by inhibiting Rac1 trafficking to the cell border. J Cell Biol 213, 385–402 (2016).

11. Wheeler, A. & Ridley, A. Why three Rho proteins? RhoA, RhoB, RhoC, and cell motility. Exp Cell Res 301, 43–49 (2004).

12. Vega, F. M., Fruhwirth, G., Ng, T. & Ridley, A. J. RhoA and RhoC have distinct roles in migration and invasion by acting through different targets. J Cell Biol 193, 655–665 (2011).

13. Ridley, A. J. RhoA, RhoB and RhoC have different roles in cancer cell migration. J Microsc 251, 242–249 (2013).

14. Dejana, E. Endothelial cell-cell junctions: happy together. Nat Rev Mol Cell Biol 5, 261–270 (2004).

15. Fernandez-Martin, L. et al. Crosstalk between reticular adherens junctions and platelet endothelial cell adhesion molecule-1 regulates endothelial barrier function. Arterioscler Thromb Vasc Biol 32, e90–e102 (2012).

16. Klomp, J. E. et al. Time-Variant SRC Kinase Activation Determines Endothelial Permeability Response. Cell Chem Biol 26, 1081–1094.e6 (2019).

17. Tzima, E. et al. A mechanosensory complex that mediates the endothelial cell response to fluid shear stress. Nature 437, 426–431 (2005).

18. Carmeliet, P. et al. Targeted Deficiency or Cytosolic Truncation of the VE-cadherin Gene in Mice Impairs VEGF-Mediated Endothelial Survival and Angiogenesis. Cell 98, 147–157 (1999).

19. Giannotta, M., Trani, M. & Dejana, E. VE-Cadherin and Endothelial Adherens Junctions: Active Guardians of Vascular Integrity. Dev Cell 26, 441–454 (2013).

20. Hayer, A. et al. Engulfed cadherin fingers are polarized junctional structures between collectively migrating endothelial cells. Nat Cell Biol 18, 1311–1323 (2016).

21. Colás-Algora, N. et al. Compensatory increase of VE-cadherin expression through ETS1 regulates endothelial barrier function in response to TNFα. Cell Mol Life Sci 77, 2125–2140 (2020).

22. Vandenbroucke, E., Mehta, D., Minshall, R. & Malik, A. B. Regulation of Endothelial Junctional Permeability. Ann N Y Acad Sci 1123, 134–145 (2008).

23. van Nieuw Amerongen, G. P., Vermeer, M. A. & van Hinsbergh, V. W. Role of RhoA and Rho kinase in lysophosphatidic acid-induced endothelial barrier dysfunction. Arterioscler Thromb Vasc Biol 20, E127–133 (2000).

24. Schmidt, G. et al. Gln 63 of Rho is deamidated by Escherichia coli cytotoxic necrotizing factor-1. Nature 387, 725–729 (1997).

25. Schlegel, N., Meir, M., Spindler, V., Germer, C.-T. & Waschke, J. Differential role of Rho GTPases in intestinal epithelial barrier regulation in vitro. J Cell Physiol 226, 1196–1203 (2011).

26. Aktories, K. Rho-modifying bacterial protein toxins. Pathog Dis 73(9), ftv091 (2015).

27. Ortega, M. C. et al. Activation of Rac1 and Rhoa preserve corneal endothelial barrier function. Invest Ophthalmol Vis Sci 57(14):6210–6222 (2016).

28. Reglero-Real, N. et al. Autophagy modulates endothelial junctions to restrain neutrophil diapedesis during inflammation. Immunity 54, 1989–2004.e9 (2021).

29. Klomp, J. E. et al. Time-Variant SRC Kinase Activation Determines Endothelial Permeability Response. Cell Chem Biol 26, 1081–1094 e6 (2019).

30. Tiruppathi, C., Malik, A. B., Del Vecchio, P. J., Keese, C. R. & Giaever, I. Electrical method for detection of endothelial cell shape change in real time: assessment of endothelial barrier function. Proc Natl Acad Sci USA 89, 7919–7923 (1992).

31. Birukov, K. G. et al. Shear stress-mediated cytoskeletal remodeling and cortactin translocation in pulmonary endothelial cells. Am J Respir Cell Mol Biol 26, 453–464 (2002).

32. Hoffmann, C., Aktories, K. & Schmidt, G. Change in Substrate Specificity of Cytotoxic Necrotizing Factor Unmasks Proteasome-independent Down-regulation of Constitutively Active RhoA. J Biol Chem 282, 10826–10832 (2007).

33. Aktories, K. & Hall, A. Botulinum ADP-ribosyltransferase C3: a new tool to study low molecular weight GTP-binding proteins. Trends Pharmacol Sci 10, 415–8 (1989).

34. Gabarin, R. S. et al. Intracellular and Extracellular Lipopolysaccharide Signaling in Sepsis: Avenues for Novel Therapeutic Strategies. J Innate Immun 13, 323–332 (2021).

35. Miles, A. A. & Miles, E. M. Vascular reactions to histamine, histamine-liberator and leukotaxine in the skin of guinea-pigs. J Physiol 118, 228–257 (1952).

36. Pronk, M. C. A., van Bezu, J. S. M., van Nieuw Amerongen, G. P., van Hinsbergh, V. W. M. & Hordijk, P. L. RhoA, RhoB and RhoC differentially regulate endothelial barrier function. Small GTPases 10, 466–484 (2019).

37. Colás-Algora, N. & Millán, J. How many cadherins do human endothelial cells express? Cell Mol Life Sci 76, 1299–1317 (2019).

38. Heissler, S. M. & Manstein, D. J. Nonmuscle myosin-2: mix and match. Cell Mol Life Sci 70, 1–21 (2013).

39. Agarwal, P. & Zaidel-Bar, R. Principles of Actomyosin Regulation In Vivo. Trends Cell Biol 29, 150–163 (2019).

40. Park, I. et al. Myosin regulatory light chains are required to maintain the stability of myosin II and cellular integrity. Biochem J 434, 171–180 (2011).

41. Oya, R. et al. Phosphorylation of MYL12 by Myosin Light Chain Kinase Regulates Cellular Shape Changes in Cochlear Hair Cells. J Assoc Res Otolaryngol 22, 425–441 (2021).

42. Totsukawa, G. et al. Distinct roles of MLCK and ROCK in the regulation of membrane protrusions and focal adhesion dynamics during cell migration of fibroblasts. J Cell Biol 164(3):427–39 (2004) .

43. Reinhard, N. R. et al. Spatiotemporal analysis of RhoA/B/C activation in primary human endothelial cells. Sci Rep 6, 25502 (2016).

44. Ridley, A. J. Rho family proteins: coordinating cell responses. Trends Cell Biol 11, 471–477 (2001).

45. Bagci, H. et al. Mapping the proximity interaction network of the Rho-family GTPases reveals signalling pathways and regulatory mechanisms. Nat Cell Biol 22, 120–134 (2020).

46. Santander-García, D. et al. A human cellular system for analyzing signaling during corneal endothelial barrier dysfunction. Exp Eye Res 153, 8–13 (2016).

47. Zawistowski, J. S., Sabouri-Ghomi, M., Danuser, G., Hahn, K. M. & Hodgson, L. A RhoC biosensor reveals differences in the activation kinetics of RhoA and RhoC in migrating cells. PLoS One 8, e79877 (2013).

48. Mahlandt, E. K. et al. Opto-RhoGEFs, an optimized optogenetic toolbox to reversibly control Rho GTPase activity on a global to subcellular scale, enabling precise control over vascular endothelial barrier strength. eLife 12, RP84364 (2023).

49. Holzner, S. et al. Phosphorylated cingulin localises GEF-H1 at tight junctions to protect vascular barriers in blood endothelial cells. J Cell Sci 134, jcs258557 (2021).

50. Khan, A. et al. Tumor necrosis factor-induced ArhGEF10 selectively activates RhoB contributing to human microvascular endothelial cell tight junction disruption. FASEB J 35(6),e21627 (2021).

51. Lang, P. et al. Protein kinase A phosphorylation of RhoA mediates the morphological and functional effects of cyclic AMP in cytotoxic lymphocytes. EMBO J 15, 510–519 (1996).

52. Birukova, A. A., Alekseeva, E., Mikaelyan, A. & Birukov, K. G. HGF attenuates thrombin-induced endothelial permeability by Tiaml-mediated activation of the Rac pathway and by Tiam1/Rac-dependent inhibition of the Rho pathway. FASEB J 21, 2776–2786 (2007).

53. Priya, R. et al. Feedback regulation through myosin II confers robustness on RhoA signalling at E-cadherin junctions. Nat Cell Biol 17, 1282–1293 (2015).

54. Kakiashvili, E. et al. GEF-H1 Mediates Tumor Necrosis Factor-α-induced Rho Activation and Myosin Phosphorylation. J Biol Chem 284, 11454–11466 (2009).

55. Nusrat, A. et al. Rho Protein Regulates Tight Junctions and Perijunctional Actin Organization in Polarized Epithelia. Proc Natl Acad Sci U S A 92, 10629–10633 (1995).

56. Sahai, E. & Marshall, C. J. ROCK and Dia have opposing effects on adherens junctions downstream of Rho. Nat Cell Biol 4, 408–415 (2002).

57. López-Posadas, R. et al. Rho-A prenylation and signaling link epithelial homeostasis to intestinal inflammation. J Clin Invest 126, 611–626 (2016).

58. Wei, J., Jiang, H. & Chen, X. Endothelial cell metabolism in sepsis. World J Emerg Med 14, 10 (2023).

59. Vincent, J.-L. Current sepsis therapeutics. eBioMedicine 86, 104318 (2022).

60. Ronco, C., Chawla, L., Husain-Syed, F. & Kellum, J. A. Rationale for sequential extracorporeal therapy (SET) in sepsis. Crit Care 27, 50 (2023).

61. Giamarellos-Bourboulis, E. J. et al. The pathophysiology of sepsis and precision-medicine-based immunotherapy. Nat Immunol 25, 19–28 (2024).

62. Aman, J. et al. Effective Treatment of Edema and Endothelial Barrier Dysfunction With Imatinib. Circulation 126, 2728–2738 (2012).

63. Aman, J. et al. Imatinib in patients with severe COVID-19: a randomised, double-blind, placebo-controlled, clinical trial. Lancet Resp Med 9, 957–968 (2021).

64. Duijvelaar, E. et al. Long-term clinical outcomes of COVID-19 patients treated with imatinib. Lancet Resp Med 10, e34–e35 (2022).

65. Juettner, V. V. et al. VE-PTP stabilizes VE-cadherin junctions and the endothelial barrier via a phosphatase-independent mechanism. J Cell Biol 218, 1725–1742 (2019).

66. Timmerman, I. et al. A local VE-cadherin and Trio-based signaling complex stabilizes endothelial junctions through Rac1. J Cell Sci 128, 3514–3514 (2015).

67. Ngok, S. P. et al. VEGF and Angiopoietin-1 exert opposing effects on cell junctions by regulating the Rho GEF Syx. J Cell Biol 199, 1103–1115 (2012).

68. Acharya, B. R. et al. A Mechanosensitive RhoA Pathway that Protects Epithelia against Acute Tensile Stress. Dev Cell 47, 439–452.e6 (2018).

69. Ngok, S. P. et al. TEM4 is a junctional Rho GEF required for cell–cell adhesion, monolayer integrity and barrier function. J Cell Sci 126, 3271–3277 (2013).

70. Zebda, N. et al. Interaction of p190RhoGAP with C-terminal Domain of p120-catenin Modulates Endothelial Cytoskeleton and Permeability. J Biol Chem 288, 18290–18299 (2013).

71. Pierce, R. W. et al. A p190BRhoGAP mutation and prolonged RhoB activation in fatal systemic capillary leak syndrome. J Exp Med 214, 3497–3505 (2017).

72. Khan, A. et al. ArhGEF12 activates Rap1A and not RhoA in human dermal microvascular endothelial cells to reduce tumor necrosis factor-induced leak. FASEB J 36(4), e22254. (2022).

73. Ren, X.-D. & Schwartz, M. A. Determination of GTP loading on Rho. in Methods Enzymol 325, 264–272.

74. Kremer, J. R., Mastronarde, D. N. & McIntosh, J. R. Computer Visualization of Three-Dimensional Image Data Using IMOD. J Struct Biol 116, 71–76 (1996).

75. Shevchenko, A., Wilm, M., Vorm, O. & Mann, M. Mass Spectrometric Sequencing of Proteins from Silver-Stained Polyacrylamide Gels. Anal Chem 68, 850–858 (1996).

76. Singer, M. et al. The Third International Consensus Definitions for Sepsis and Septic Shock (Sepsis-3). JAMA 315, 801–810 (2016).

