## Supplemental Information for "Harnessing homeostatically active RhoC at cell junctions preserves human endothelial barrier function during inflammation"

Natalia Colás-Algora et al.

9 Extended Data Figures

1 Extended Data Table

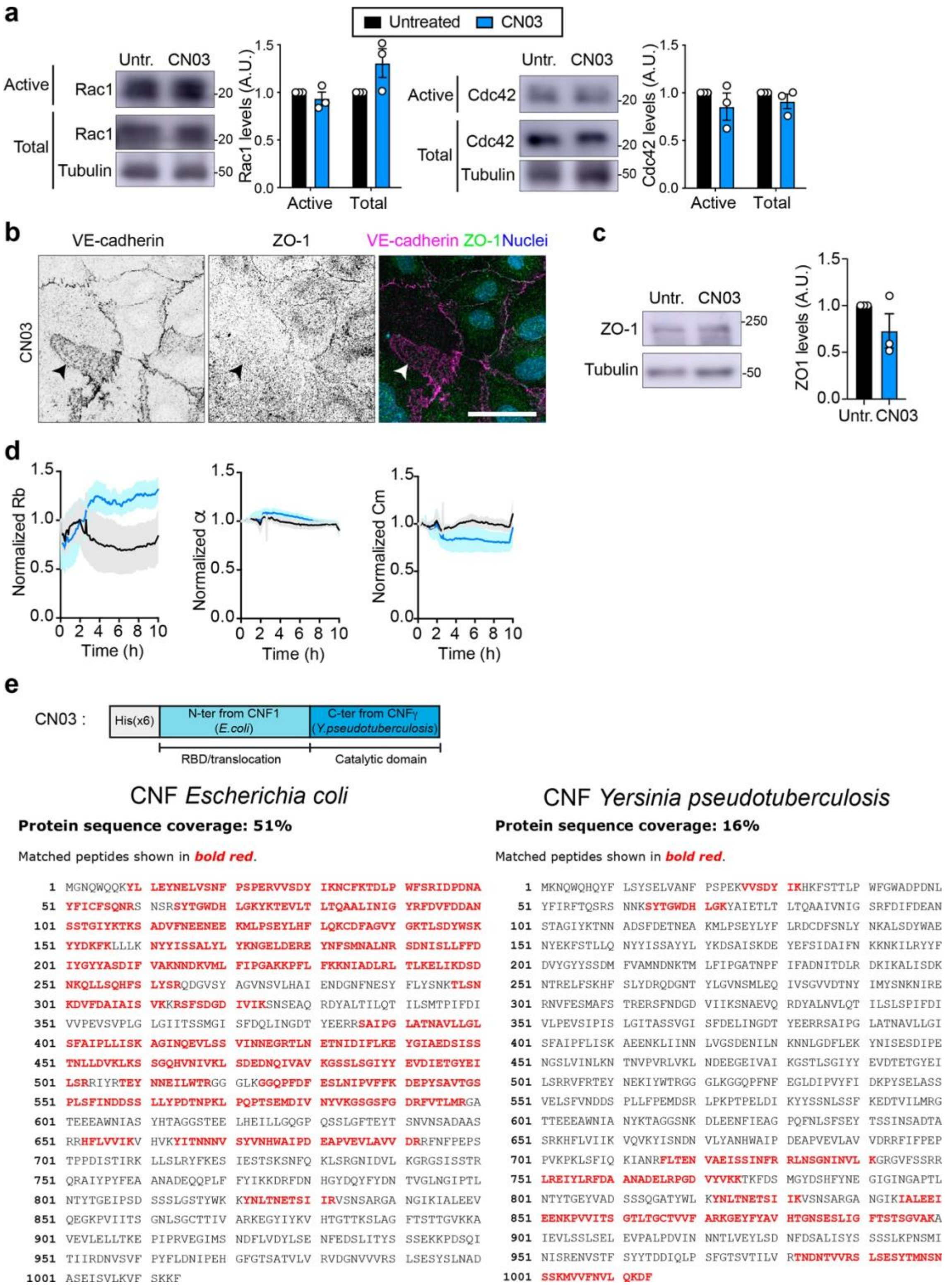


**Extended Data Figure 1. The RhoA subfamily activator CN03 does not modulate the activity of Rac1 and Cdc42. Related to Figure 1. a,** Confluent HUVECs were exposed or not to 1.5 μg/ml CN03 for 3 h, lysed and the indicated active GTPases were detected by pull-down assays as in Figure 1a. **b,** Confluent HUVECs were exposed to 1.5 μg/ml for 3 h, fixed and stained for VE-cadherin and ZO-1. Arrowheads point at reticular AJs devoid of ZO-1. Nuclei were visualized with DAPI. **c,** Confluent HUVECs were exposed or not to 1.5 μg/ml CN03 for 3 h, lysed and blotted for ZO-1 and tubulin as a loading control. Right plots show quantification of ZO-1 expression levels form three independent experiments. **d,** Rb, $\alpha$ and Cm values corresponding to the TEER analyses of Figure 1g. **e,** The mass spectrometry analysis of CN03 identified the indicated peptides in red, which demonstrates that this chimeric protein is composed of the N-terminal domain of the CNF from *Escherichia coli* and the C-terminal, catalytic domain, corresponding to the CNF from *Yersinia pseudotuberculosis*.

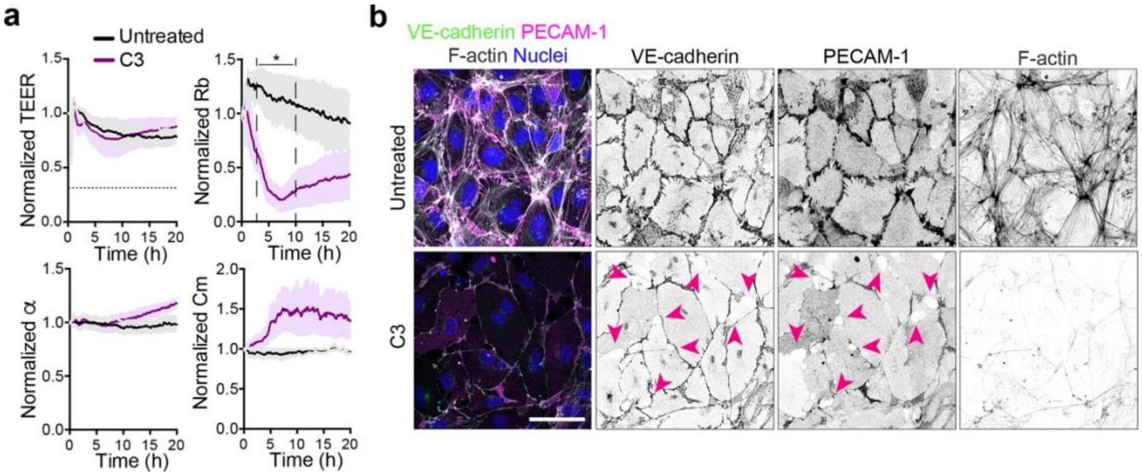


**Extended Data Figure 2. The RhoA subfamily inhibitor C3 transferase increases endothelial paracellular permeability. a,** HUVECs were cultured at confluency on ECIS arrays for 72 h, exposed or not to 1μg/ml of C3 transferase, and TEER measured for 20 h. Rb, α and Cm were calculated by the ECIS software. Graphs show the mean ± SEM from at least three independent experiments. *, $P<0.05$. **b,** HUVECs were cultured at confluency for 72 h on glass coverslips, exposed or not to 1μg/ml of C3 transferase for 20 h, fixed, stained for VE-cadherin, PECAM-1 and F-actin and analyzed by confocal microscopy. Purple arrowheads point at intercellular gaps induced by C3. Nuclei were visualized with DAPI. Scale bar, 50 μm.

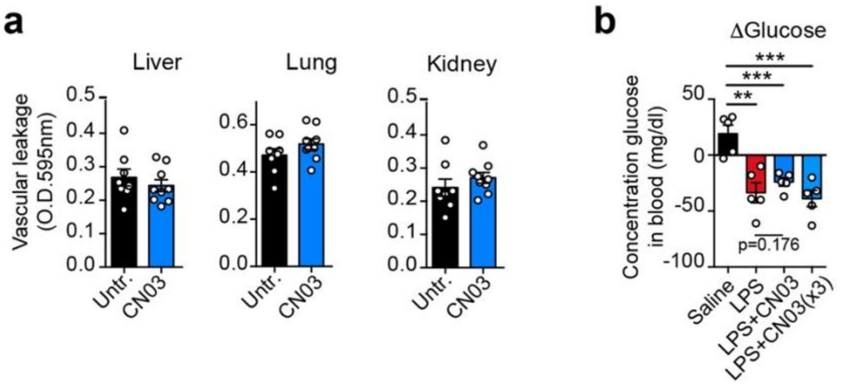


**Extended Data Figure 3. Effect of local intradermal injection of CN03 in the microvascular permeability of the indicated tissues. Effect of the intraperitoneal injection of CN03 and LPS in blood glucose levels. Related to Figure 4. a,** Local inflammation in skin does not alter systemic microvascular permeability. Miles assay of the indicated organs from experiments of local inflammatory challenge in murine skin shown in Figure 4a. **b,** Related to Figure 4b. Mice were intra-peritoneally injected with saline, LPS+saline, LPS + 0.175 mg/kg CN03 and LPS + 0.525 mg/kg (CN03x3) for 3.5 h. Blood was collected before and after LPS injection and the differences represented in the graph. Graphs show the mean ± SEM from at least five animals per condition. **, P<0.001; ***, P<0.0001.

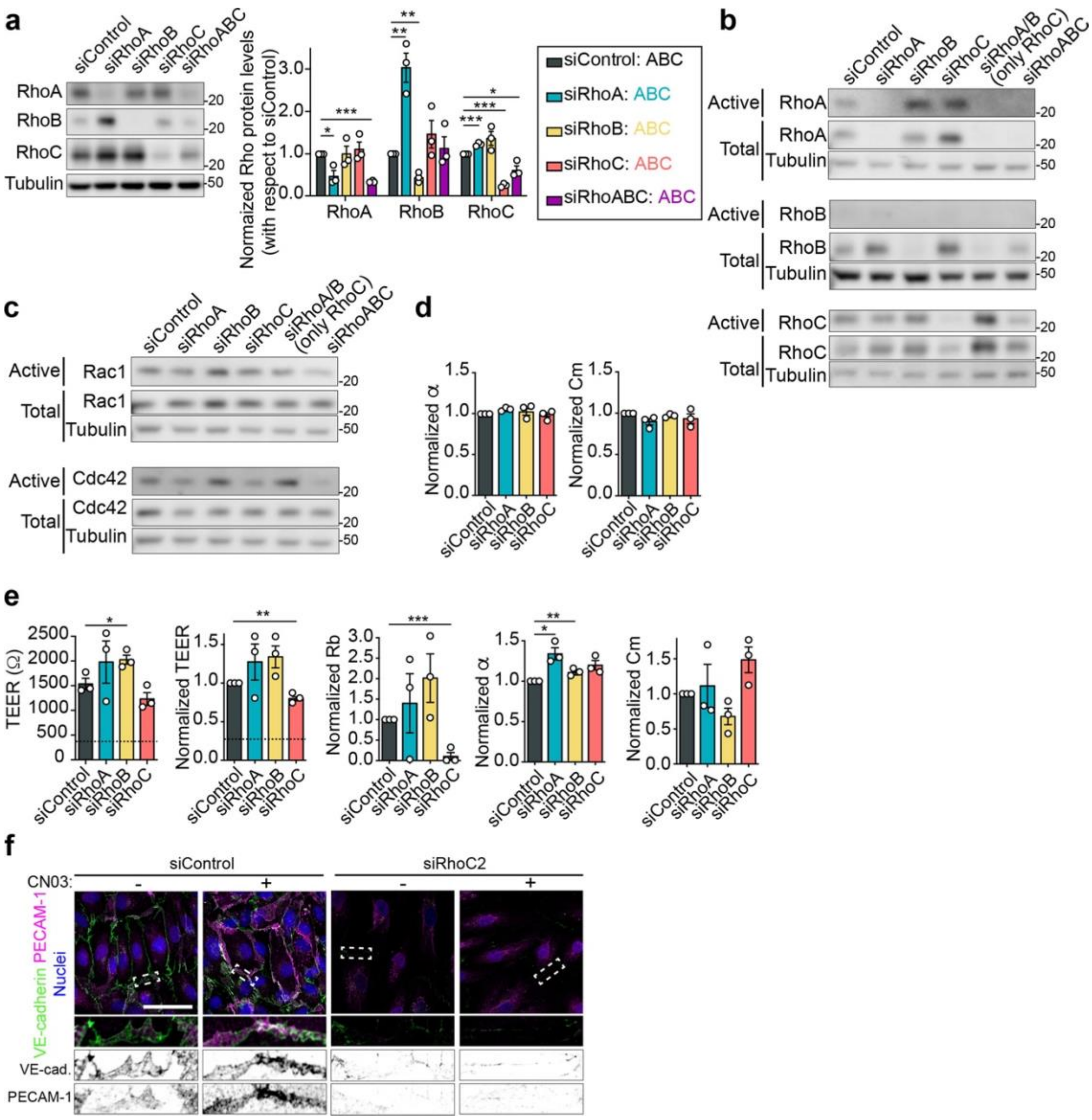


**Extended Data Figure 4. RhoC mediates the protective effect of CN03 on microvascular endothelial barrier function. Related to Figure 5. a,** Compensatory changes of expression of RhoA subfamily members upon single knockdown of RhoA, RhoB and RhoC in HDMVEC. **b,** The compensatory changes of Rho expression observed during single or double knockout of RhoA, RhoB and RhoC correlate with changes in their activity, measured by pull-down assays with GST conjugated with the RBD of rothekin (Active). **c,** Effect of gene silencing of the RhoA subfamily members on the activity of Rac1 and Cdc42, measured by pull-down assays with GST conjugated to the Rac/Cdc42 binding domain of PAK. **d,** α and Cm values corresponding to the experiments shown in Figure 5b in HUVECs. **e,** Specific role of RhoC in maintaining human microvascular endothelial barrier function and regulating paracellular permeability. Absolute and normalized TEER, Rb, α and Cm values of HDMVEC transfected with the indicated siRNA for 72 h and subjected to ECIS analysis. Plots show the mean ± SEM from at least three independent experiments. **f,** Confocal images of HUVECs transfected with siRhoC2 in parallel to the transfections performed for Figure

5c. Nuclei were visualized with DAPI. Quantifications are included in Figure 5d. *, P< 0.05; **, P< 0.01; *** P<0.001.

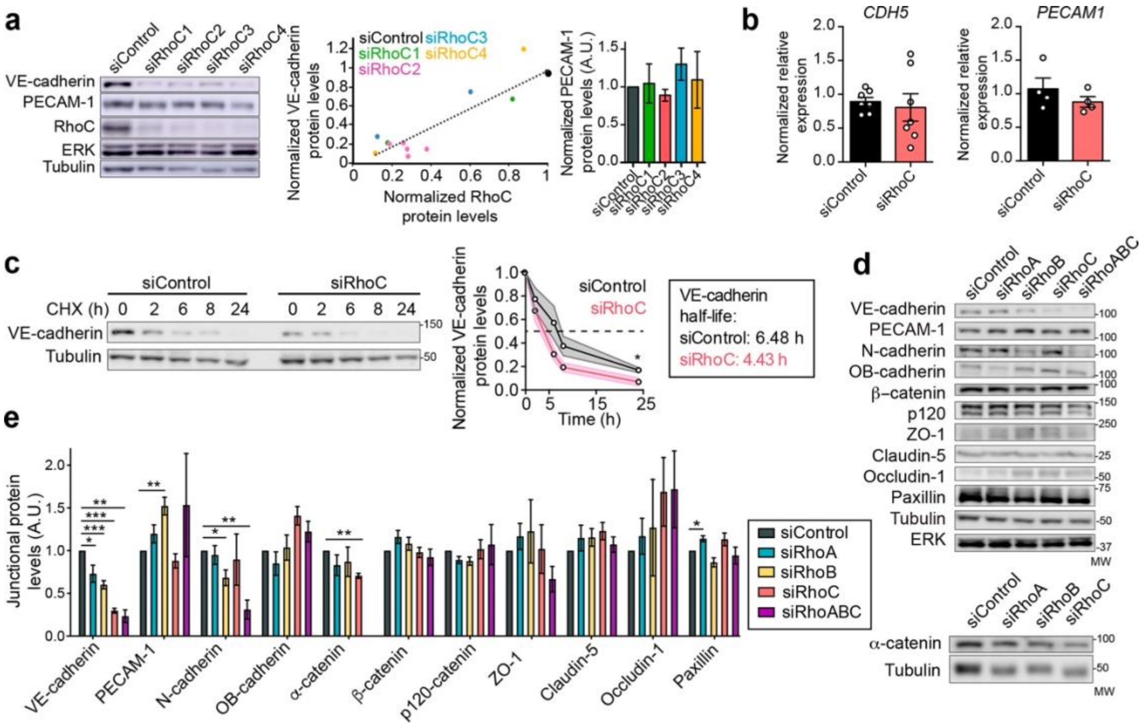


**Extended Data Figure 5. Effect of single knockout of RhoA subfamily members on the expression of endothelial junctional proteins. a,** Effect of single transfection of four different RhoC siRNA oligonucleotides on VE-cadherin and PECAM-1 expression levels. HUVECs transfected with the indicated siRNA for 72 h were lysed and the indicated proteins detected by western blot (left). Tubulin and ERK were immunoblotted as loading controls. Central plot shows the correlation between RhoC and VE-cadherin expression levels. Right plot shows the quantification of PECAM-1 in siRNA-transfected cells. (A.U.) arbitrary units. Plots show the mean + SEM from three independent experiments. **b,** qPCR analyses of cells transfected with the indicated siRNAs for 72 h. Plots show the mean + SEM from seven independent experiments. **c,** HUVECs were transfected with the indicated siRNAs for 72 h and incubated with 25 μg/ml of the protein synthesis inhibitor cycloheximide (CHX) for the indicated time periods, lysed and immunoblotted for the indicated proteins. Tubulin was blotted as a control of a protein with longer half-life. Right graph shows the quantification of VE-cadherin protein expression levels normalized to those detected at t=0 of CHX incubation and represents the mean + SEM from three independent experiments. **d,** HUVECs were transfected with the indicated siRNAs for 72 h, lysed and immunoblotted for the indicated junctional proteins. Tubulin and ERK were blotted as loading controls. **e,** Quantification of junctional protein levels detected by western blot and normalized to values in siControl cells. Plots show the mean ± SEM from at least three independent experiments *, P< 0.05; **, P< 0.01; *** P<0.001.

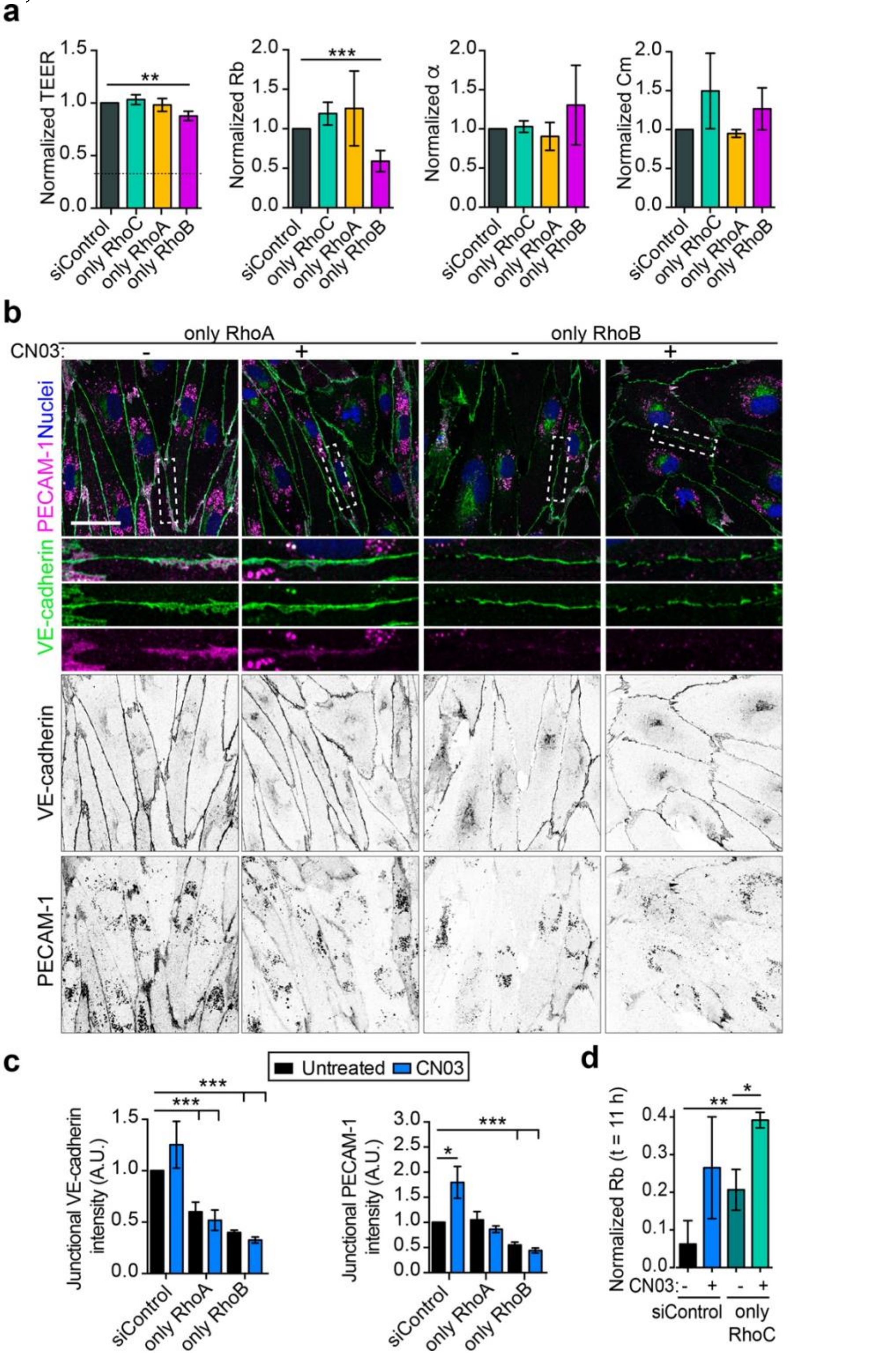


**Extended Data Figure 6. Effect of double knockdown of RhoA subfamily members on endothelial barrier function. Related to Figure 5. a,** TEER analyses and subsequent Rb, α and Cm calculations of only RhoA, only RhoB and only RhoC HUVECs. **b,** Related to Figure 5f. VE-cadherin and PECAM-1 distribution in only RhoA and only RhoB cells. Effect of 3 h exposure to CN03. Nuclei were visualized with DAPI. Scale bar, 50 μm. **c,** Quantification of junctional VE-cadherin and PECAM-1 intensities. **d,** Quantification of

Rb values 11 h post-LPS stimulation in experiment of Figure 5g. Graphs show the mean ± SEM from three independent experiments *, P< 0.05; **, P< 0.01; *** P<0.001.

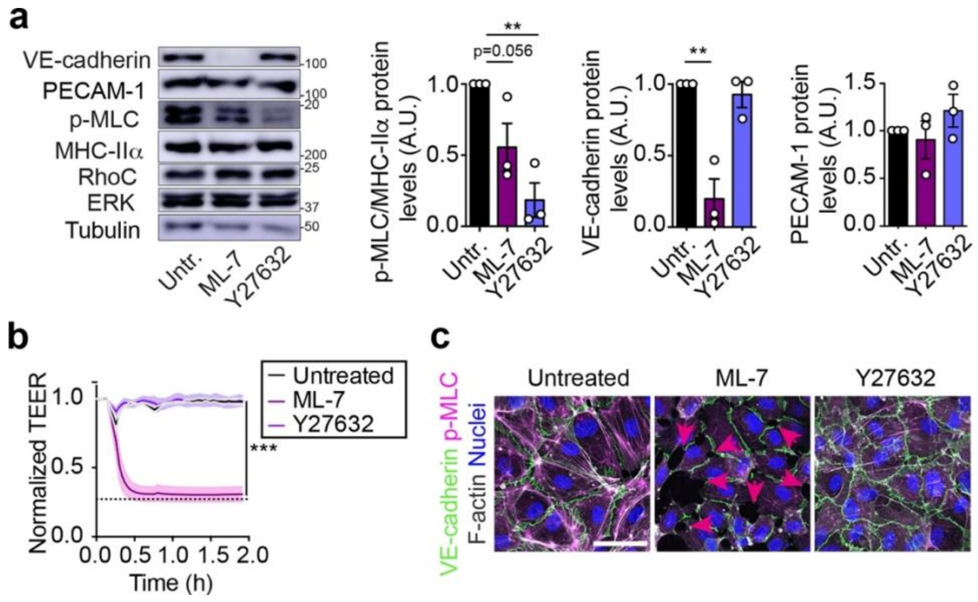


**Extended Data Figure 7. MLCK inhibition, but not ROCK inhibition, reduces VE-cadherin expression and disrupt endothelial barrier function. a,** HUVECs were cultured at confluence for 72 h, left untreated or incubated with 50 μM ML-7 or 5 μM Y-27632 for 2 h, lysed and immunoblotted for the indicated proteins. Tubulin and ERK were blotted as loading controls. Plots show quantification of expression for the indicated proteins from three independent experiments. Mean ± SEM is shown. **, P< 0.01. **b,** HUVECs were cultured at confluence for 72 h on ECIS arrays, left untreated or incubated with 50 μM ML-7 or μM Y-27632 for 2 h. ***, P< 0.001. **c,** HUVECs were cultured at confluence for 72 h, left untreated or incubated with 50 μM ML-7 or 5 μM Y-27632 for 2 h, fixed, immunostained for the indicated proteins and analyzed by confocal microscopy. Arrowheads point at intercellular gaps. Nuclei were visualized with DAPI. Scale bar, 50 μm.

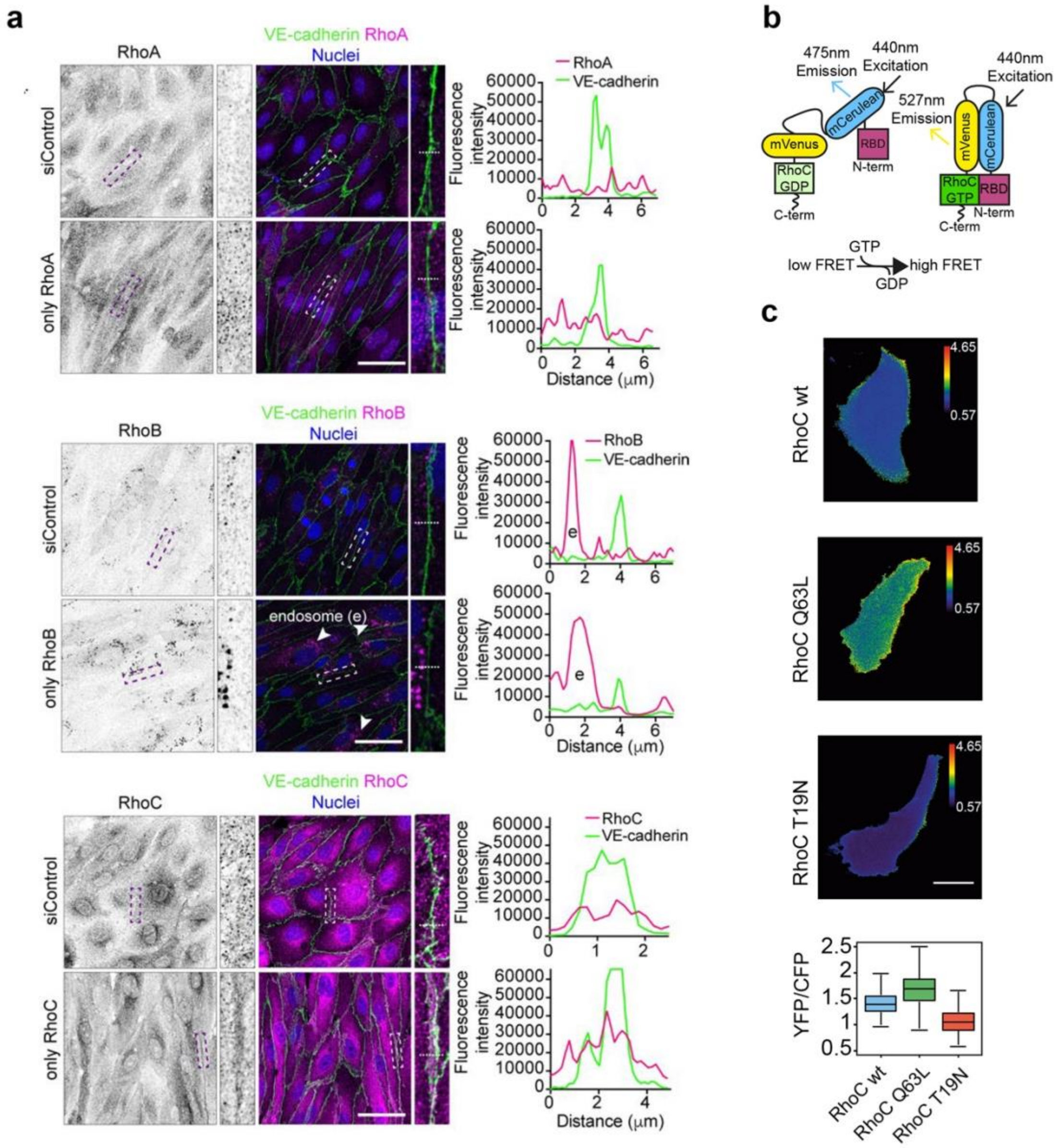


**Extended Data Figure 8. Confocal analysis of RhoA, RhoB and RhoC in siControl, only RhoA, only RhoB and only RhoC HUVECs. Related to Figure 7. a,** Cells were cultured at confluency for 72 h, fixed and stained for VE-cadherin and RhoA, RhoB or RhoC with specific antibodies when indicated. Nuclei were stained with DAPI. Lateral images show a three-fold enlargement of the squared areas. Right graphs show the intensity profiles of the indicated stainings in the discontinuous lines from the enlarged images. Scale bar, 50 μm. **b,** Conformational changes of RhoC-FLARE chimeric protein in response to RhoC activation, taken from (Zawistowski et al., 2013). **c,** Constitutively active RhoC Q63L FLARE shows higher FRET signal (YFP/CFP ratio) than RhoC-FLARE. Dominant negative RhoC T19N FLARE has lower FRET signal than RhoC-FLARE and RhoC Q63L FLARE.

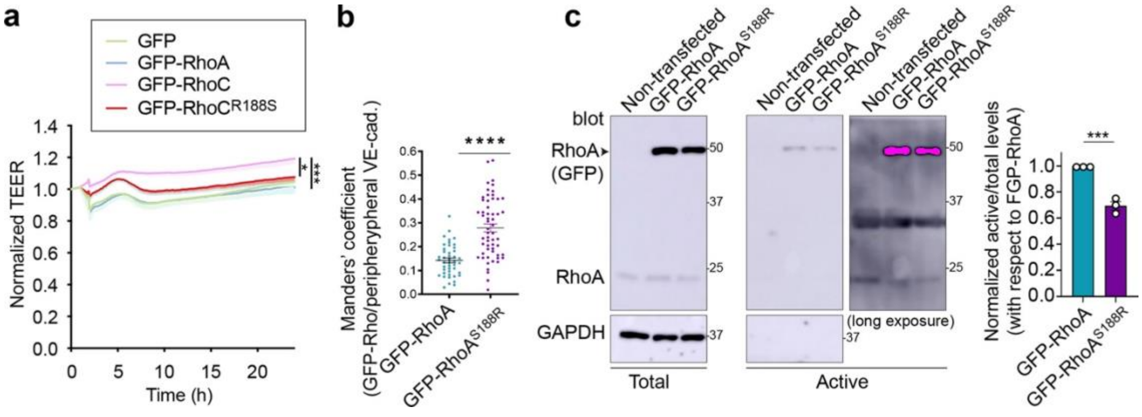


**Extended Data Figure 9. The pro-barrier role of RhoC is regulated by a single basic residue in the hypervariable region, which is not present in RhoA. Related to Figure 8. A,** TEER values of experiments in Figure 8d. **b**,**c,** Substitution of S188 in the hypervariable region of GFP-RhoA by R alters its junctional localization but does not increase its activity in homeostatic conditions. **a**, Expression plasmids containing the indicated chimeras were ectopically expressed for 24 h in HUVECs, cells were fixed, stained for VE-cadherin and junctional co-localization quantified with Manders' coefficient. **c,** Confluent HUVECs expressing the indicated GFP-Rho proteins for 48 h were lysed and subjected to pull-down assays with GST-RTK to detect GTP-loaded RhoA. Total and active fractions were immunoblotted with the indicated antibodies. Right graph shows the quantification of GFP-RhoA$^{S188R}$ activation relative to that of GFP-RhoA. Signal-saturated, long exposure is shown to visualize the band of endogenous RhoA. Plots show the mean ± SEM from three independent experiments. *, P< 0.05; *** P<0.001; **** P<0.0001.

**Extended Data Table 1.** Raw data of relative protein abundance for junctional and actomyosin-related proteins for each triplicate of siControl, siRhoC1 and siRhoC2 cells.

| Accession | Description | Sum PEP Score | # PSMs | # Peptides | # Unique Peptides | Abundances (Normalized): siControl | | | Abundances (Normalized): siRhoC | | | | | |
|---|---|---|---|---|---|---|---|---|---|---|---|---|---|---|
| | | | | | | siControl (F1) | siControl (F2) | siControl (F3) | siRhoC1 (F4) | siRhoC2 (F7) | siRhoC1 (F5) | siRhoC2 (F8) | siRhoC1 (F6) | siRhoC2 (F9) |
| JUNCTIONAL-RELATED PROTEINS | | | | | | | | | | | | | | |
| P35221 | Catenin alpha-1 OS=Homo sapiens OX=9606 GN=CTNNA1 PE=1 SV=1 | 801.582 | 840 | 56 | 55 | 559985442.3 | 561130485.5 | 520936351.3 | 382149934.6 | 403008717.7 | 401003762 | 381571611.8 | 368738296.2 | 375213282.4 |
| Q9UI47 | Catenin alpha-3 OS=Homo sapiens OX=9606 GN=CTNNA3 PE=1 SV=2 | 8.591 | 27 | 3 | 2 | 4970552.278 | 3853573.886 | 4162040.12 | 2031612.22 | 4701527.824 | 3720895.25 | 4641260.104 | 2665957.636 | 5092961.456 |
| O60716 | Catenin delta-1 OS=Homo sapiens OX=9606 GN=CTNND1 PE=1 SV=1 | 459.8 | 433 | 43 | 43 | 214673612 | 216780752.3 | 215325664.3 | 208506568.3 | 155429562.1 | 216982366.8 | 154022988.8 | 214604941.8 | 153550223.6 |
| P35222 | Catenin beta-1 OS=Homo sapiens OX=9606 GN=CTNNB1 PE=1 SV=1 | 298.844 | 353 | 28 | 24 | 179062831 | 175100989.4 | 165827670.9 | 228101187.6 | 134857224.8 | 241571944.7 | 133719676.6 | 228199162.6 | 131448570.2 |
| Q8WYA6 | Beta-catenin-like protein 1 OS=Homo sapiens OX=9606 GN=CTNNBL1 PE=1 SV=1 | 99.386 | 98 | 10 | 10 | 15328145.98 | 14477173.64 | 16232851.42 | 11798721.4 | 14839870.66 | 12936250.56 | 18629648.52 | 10673180.08 | 19961094.97 |
| Q07157 | Tight junction protein ZO-1 OS=Homo sapiens OX=9606 GN=TJP1 PE=1 SV=3 | 233.531 | 274 | 25 | 25 | 58972291.86 | 53756660.85 | 57032672.65 | 53663364.92 | 41371650.25 | 50702683.05 | 35812538.58 | 50634542.36 | 40393815.81 |
| Q9UDY2 | Tight junction protein ZO-2 OS=Homo sapiens OX=9606 GN=TJP2 PE=1 SV=2 | 364.851 | 453 | 33 | 33 | 192941587.6 | 187273981 | 211911490.1 | 168264563.7 | 130359370.8 | 204075848.7 | 164641744.6 | 209905095.6 | 176734039.8 |
| P17302 | Gap junction alpha-1 protein OS=Homo sapiens OX=9606 GN=GJA1 PE=1 SV=2 | 26.89 | 15 | 6 | 6 | 2263900.978 | 2411541.473 | 2316323.319 | 5302322.867 | 4727746.633 | 5829436.5 | 4015626.249 | 4005989.365 | 4485396.253 |
| Q9Y624 | Junctional adhesion molecule A OS=Homo sapiens OX=9606 GN=F11R PE=1 SV=1 | 67.424 | 123 | 7 | 7 | 32594089.19 | 30461747.07 | 33040743.46 | 49943420.43 | 25000050.67 | 49286072.56 | 23351438.21 | 51548505.77 | 22464269.3 |
| Q9BX67 | Junctional adhesion molecule C OS=Homo sapiens OX=9606 GN=JAM3 PE=1 SV=1 | 9.523 | 16 | 2 | 2 | 4804274.957 | 5728182.315 | 4283642.797 | 8339583.512 | 2747951.257 | 6743229.125 | 2878039.853 | 8303184.709 | 3009787.953 |
| Q5JTD0 | Tight junction-associated protein 1 OS=Homo sapiens OX=9606 GN=TJAP1 PE=1 SV=1 | 13.818 | 5 | 2 | 2 | 447887.3169 | 822411.5429 | 756627.4723 | 764616.6371 | | 1220157.406 | 626500.4133 | 1171662.656 | 655571.3365 |
| P14923 | Junction plakoglobin OS=Homo sapiens OX=9606 GN=JUP PE=1 SV=3 | 390.959 | 605 | 33 | 29 | 264186463.3 | 270005771.2 | 266426832.1 | 219274025.6 | 99350915.28 | 210708022.5 | 104101199 | 205834991.1 | 99454735.38 |
| P16284 | Platelet endothelial cell adhesion molecule OS=Homo sapiens OX=9606 GN=PECAM1 PE=1 SV=2 | 677.943 | 869 | 43 | 43 | 495979664.5 | 434788086.4 | 433101295.3 | 1094939209 | 398963005.9 | 1020042283 | 372665900.2 | 1056644896 | 341426537.7 |
| Q9P266 | Junctional protein associated with coronary artery disease OS=Homo sapiens OX=9606 GN=JCAD PE=1 SV=3 | 36.53 | 43 | 6 | 6 | 5700988.693 | 5764526.237 | 4475573.19 | 7844127.81 | 3505463.095 | 4559708.063 | 3707000.421 | 4799689.813 | 9345004.767 |
| P33151 | Cadherin-5 OS=Homo sapiens OX=9606 GN=CDH5 PE=1 SV=5 | 157.527 | 154 | 15 | 15 | 127071639.2 | 121854529.6 | 118365264.7 | 145584561.9 | 61800058.91 | 140576172.1 | 54917389.63 | 141795486.1 | 55036578.42 |
| ACTIN-RELATED PROTEINS | | | | | | | | | | | | | | |
| P60660 | Myosin light polypeptide 6 OS=Homo sapiens OX=9606 GN=MYL6 PE=1 SV=2 | 292.564 | 1212 | 16 | 13 | 1238104992 | 1308745618 | 1206879583 | 736373142.1 | 1085152421 | 796489122.7 | 1044736779 | 756168620.4 | 1015424357 |
| P14649 | Myosin light chain 6B OS=Homo sapiens OX=9606 GN=MYL6B PE=1 SV=1 | 55 | 256 | 4 | 1 | 1681217.192 | 2117557.719 | 2133929.937 | 551438.2306 | 1219591.06 | 950138.5625 | 1346495.938 | 733836.3389 | 1270712.543 |
| P24844 | Myosin regulatory light polypeptide 9 OS=Homo sapiens OX=9606 GN=MYL9 PE=1 SV=4 | 50.427 | 53 | 4 | 1 | 7510293.434 | 8885051.824 | 8657377.651 | 2313868.105 | 4933578.135 | | | 1475795.305 | 4375801.334 |
| O14950 | Myosin regulatory light chain 12B OS=Homo sapiens OX=9606 GN=MYL12B PE=1 SV=2 | 128.822 | 152 | 11 | 8 | 252516431.7 | 278248465.8 | 261750878.9 | 139877441.1 | 204185895.9 | 163783372.2 | 203008089.9 | 161255028.5 | 202763587.6 |
| P63261 | Actin, cytoplasmic 2 OS=Homo sapiens OX=9606 GN=ACTG1 PE=1 SV=1 | 1738.412 | 11421 | 60 | 3 | 450681772.1 | 403319929.2 | 486915882.7 | 556963490.4 | 426364569.8 | 566002357.4 | 418960275 | 599404362.6 | 480969183.7 |
